# Black Dots: Microcontact-Printed, Reference-Free Traction Force Microscopy

**DOI:** 10.1101/2021.08.02.454500

**Authors:** Kevin M. Beussman, Molly Y. Mollica, Andrea Leonard, Jeffrey Miles, John Hocter, Zizhen Song, Moritz Stolla, Sangyoon J. Han, Ashley Emery, Wendy E. Thomas, Nathan J. Sniadecki

**Affiliations:** Department of Mechanical Engineering, University of Washington, Seattle, WA; Department of Bioengineering, University of Washington, Seattle, WA; Bloodworks Northwest Research Institute, Seattle, WA; Department of Biostatistics, University of Washington, Seattle, WA; School of Computer Science & Engineering, University of Washington, Seattle, WA; Division of Hematology, Department of Medicine, University of Washington, Seattle, WA; Department of Biomedical Engineering, Michigan Technological University, Houghton, MI; Institute for Stem Cell and Regenerative Medicine, University of Washington, Seattle, WA; Department of Laboratory Medicine & Pathology, University of Washington, Seattle, WA; Resuscitation Engineering Science Unit (RESCU), University of Washington, Seattle, WA

**Keywords:** Microcontact printing, Traction force microscopy, Cell mechanics, Platelets, Polydimethylsiloxane (PDMS)

## Abstract

Measuring the traction forces produced by cells provides insight into their behavior and physiological function. Here, we developed a technique (dubbed ‘black dots’) that microcontact prints a fluorescent micropattern onto a flexible substrate to measure cellular traction forces without constraining cell shape or needing to detach the cells. To demonstrate our technique, we assessed human platelets, which can generate a large range of forces within a population. We find platelets that exert more force have more spread area, are more circular, and have more uniformly distributed F-actin filaments. As a result of the high yield of data obtainable by this technique, we were able to evaluate multivariate mixed effects models with interaction terms and conduct a clustering analysis to identify clusters within our data. These statistical techniques demonstrated a complex relationship between spread area, circularity, F-actin dispersion, and platelet force, including cooperative effects that significantly associate with platelet traction forces.

**GRAPHICAL ABSTRACT:** 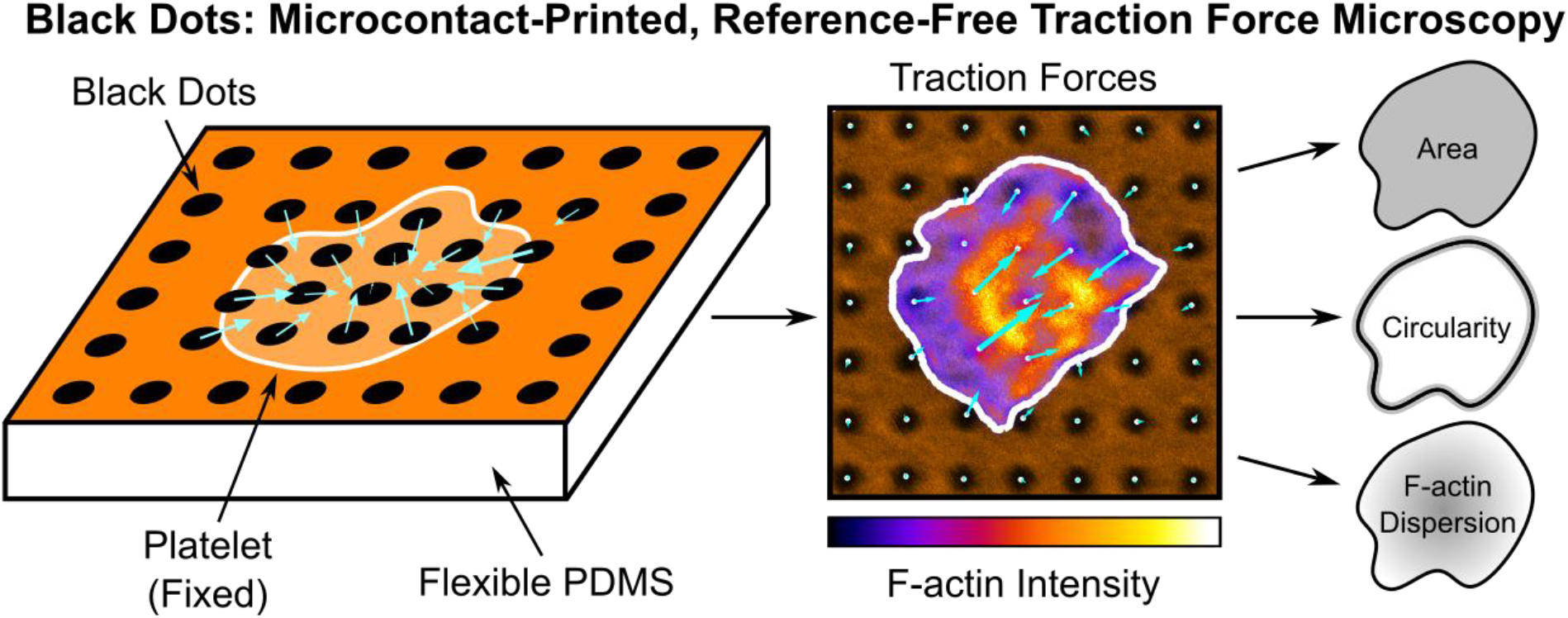

## 1. INTRODUCTION

Cells use forces to migrate, contract, and probe their physical environment[1,2]. These forces arise from interactions of cytoskeletal proteins, which transmit cellular forces to the extracellular matrix. The cellular forces can ultimately cause deformation of the surrounding environment. By measuring the deformation of the underlying substrate, cellular forces can be estimated. This principle has been used to develop techniques such as membrane wrinkling, traction force microscopy, and microposts, among many others to measure single-cell forces[3–6]. However, existing methods have several drawbacks including the limited number of cells that can be measured per experiment or inadvertent impact on cell functions by strictly constraining cell size and shape.

Traction force microscopy (TFM) is one of the most widely used techniques for measuring forces from single cells. In TFM, cellular forces are determined from the displacement of fluorescent particles embedded within a flexible substrate[7–9]. Often, a pair or series of images are required to track a cell’s forces: a reference image of the undeformed substrate and one or more images of the displacements caused by the cells. For this reason, TFM is a relatively low-yield assay and is incompatible with immunofluorescent staining. To side-step the requirement of multiple images, reference-free TFM approaches have been developed where markers are fabricated on a substrate in a pattern instead of being distributed randomly[10]. Since only a single image is required for the measurement of traction forces, reference-free TFM is compatible with fixed samples and immunofluorescent staining because the cells do not need to be detached. While reference-free TFM can increase the number of cells that can be efficiently analyzed, many of the existing methods provide a large degree of constraint on the adhesion and spreading of a cell, impacting their physiological significance[11–14].

Platelets use their traction forces to adhere and form a hemostatic plug that stops bleeding[15–17]. During this process, the actin cytoskeleton of a platelet drives its shape change, spreading, and production of traction forces. Measuring these forces for individual platelets is challenging due to their small size (2-5 μm in diameter)[18], their ability to produce strong forces[19], and their sensitivity to collection and handling techniques[20]. It has been shown that the spread area of platelets correlated with the overall magnitude of their traction forces[21,22]. While time-dependent changes in platelet shape and cytoskeletal structure have also been observed[23,24], it is not known how these factors impact traction forces in platelets. Moreover, these factors may be interrelated because the actin cytoskeleton underlies changes in shape and spreading. Previous measurements of platelet forces have used atomic force microscopy[25], classical TFM[19,21,22], reference-free TFM[12], and nanoposts[26] to elucidate properties of single platelets such as their temporal and directional contraction dynamics, the function of platelet mechanoreceptors, and the influence of biochemical and mechanical cues on platelet contraction. While these existing methods have allowed for understanding of important biophysical properties of platelets, they have been hampered by constraints on the shape or spreading of platelets and/or their low yield, often analyzing fewer than thirty platelets per condition.

Here, we present a microcontact-printed, reference-free TFM technique for measuring single-cell forces without constraining cell shape and size. Our method relies on microcontact printing to deposit a grid of fluorescently labeled bovine serum albumin (BSA) onto a flexible polydimethylsiloxane (PDMS) substrate. This procedure results in a fluorescent surface with a pattern of circular islands that are non-fluorescent (Fig. 1A), hence the technique is termed ‘black dots.’ The black dots technique offers several advantages over existing methods for measuring single-cell forces: 1) it is high-yield due to the ability to measure force with a single image, 2) it is compatible with immunofluorescent staining so that traction forces can be measured alongside analysis of structure and/or localization, and 3) it does not constrain cell shape and size due to the substrate containing a contiguous adhesive protein. With this approach, we characterized forces, cytoskeletal structures, and geometric properties of more than five hundred human platelets for linear mixed-effects modeling and K-means clustering, from which we identified that platelet size, shape, and cytoskeletal structure have both independent and cooperative contributions to platelet force.

**Figure 1.**
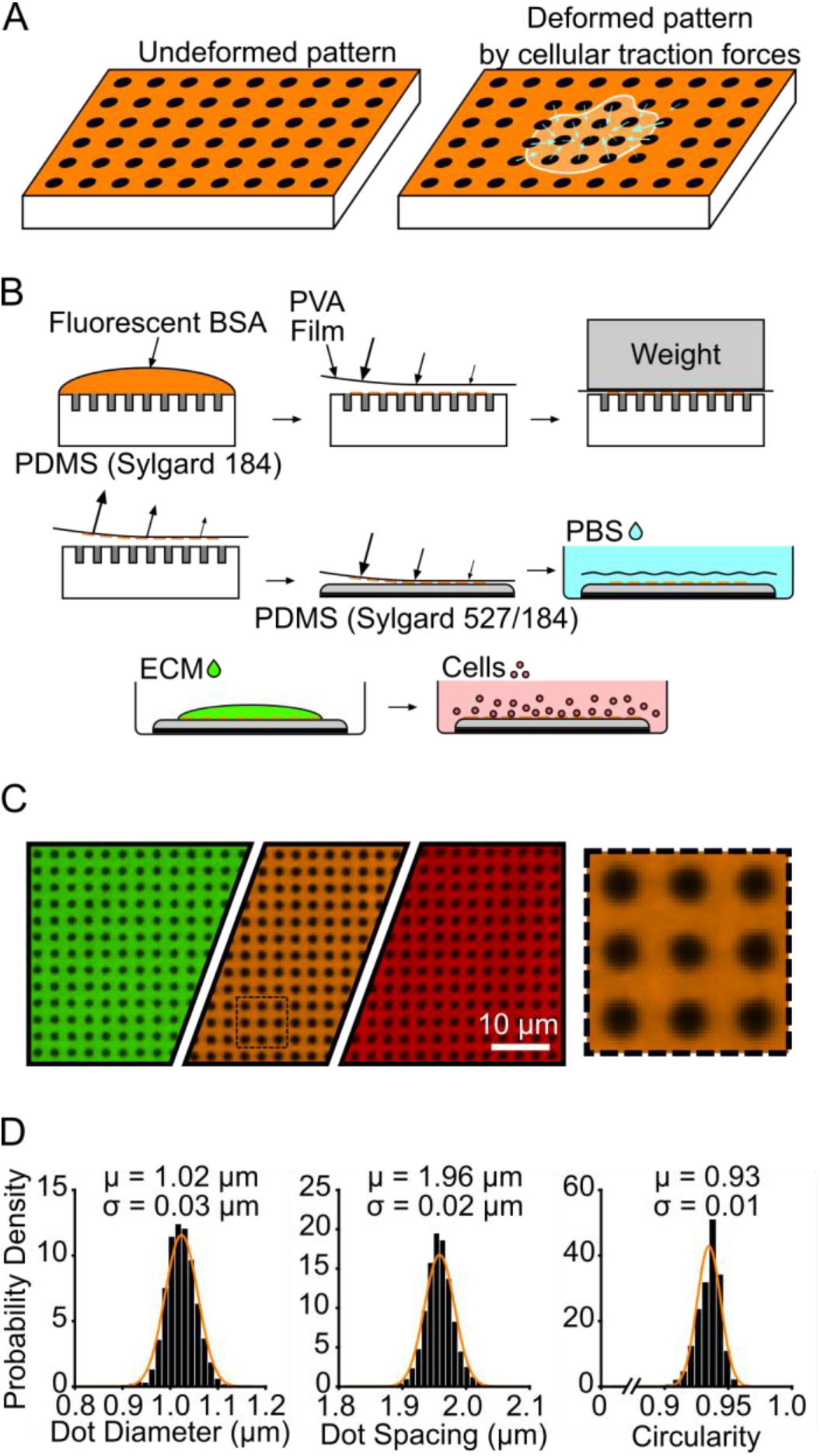
Black dots overview, manufacturing, and characterization. (A) Principle of black dots, where tension from an adhered cell causes the pattern of dots to displace. (B) Manufacturing black dots substrates using microcontact printing and a sacrificial PVA film to transfer an array of fiducial markers from a patterned stamp to a soft substrate. (C) Example of final manufactured substrate that can be made in the desired fluorescent channel using different fluorescent BSA such as BSA-Alexa Fluor 488 (green), BSA-Alexa Fluor 594 (orange), and BSA-Alexa Fluor 647 (red). The black dotted line area is shown on the right, scaled up 4X larger. (D) Characterization of diameter, center-center spacing, and circularity of black dots. μ = mean, σ = standard deviation. Data from 25,081 individual dots from 2 substrates. Y-axis is Probability Density for all three plots. Normal Gaussian probability density functions are overlayed.

## 2. MATERIALS AND METHODS

### 2.1. Microfabrication of patterned stamp

First, a silicon master mold with an array of vertical pillars was created with the desired pattern size by photolithography as described previously[26]. Briefly, photoresist was spun onto a silicon wafer and an e-beam lithography system was used to pattern circles of the desired diameter and center-to-center spacing. The photoresist was then developed and etched to create a master containing an array of vertical silicon pillars. For cells as small as platelets, we used a pattern with a diameter of 850 nm and center-to-center spacing of 2 μm, and the final etched pattern had a height of 3.5 μm. Larger cell types are amenable to larger pattern sizes which can be easier to image.

To generate the stamps for patterning the fluorescent protein, PDMS (Sylgard 184, Dow Corning) at a 10:1 base to curing agent ratio is poured onto the master mold and cured in a 110 °C oven for 20 minutes. The cured PDMS is peeled from the master mold, revealing a negative version of the original pattern: a grid of holes instead of pillars. Edges of the stamp were trimmed with sharp razors and stored in enclosed petri dishes prior to use.

### 2.2 Sacrificial PVA film production

Poly(vinyl alcohol) (PVA) films for transferring the fluorescent pattern were made following previously described protocols with some modifications[27,28]. A mixture of 0.55 g PVA powder (Sigma) was mixed with 15 mL DI water and heated for 30 minutes at 110 °C until the powder fully dissolved. A standard 10 cm Petri dish was plasma treated for 10 seconds to help the final film remain attached to the dish. After cooling down to room temperature, the liquid PVA mixture was poured into the plasma-treated dish. The dish was left uncovered in a 65 °C oven overnight to allow the liquid to completely evaporate. The next day, the dish was removed from the oven revealing a thin, dried PVA film loosely attached to the bottom of the Petri dish. The film was cut into appropriate sized pieces and used as needed, or the dish was covered and sealed with parafilm for longer term storage.

### 2.3. Flexible PDMS substrate preparation

Flexible 13.5 kPa substrates were manufactured as previously published[29,30]. Soft PDMS (Sylgard 527, 1:1 ratio of parts A and B, Dow Corning) and normal PDMS (Sylgard 184, 10:1 ratio of base to curing agent) were first prepared separately and allowed to degas for at least 20 minutes under vacuum. The two types of PDMS were then mixed to form a mixture of 5% Sylgard 184 and 95% Sylgard 527 by weight, and the mixture was degassed for 20 minutes under vacuum. Round glass coverslips (25 mm diameter, #1 thickness, VWR) were plasma treated for 30 seconds (Plasma Prep II, SPI Supplies) and a 100-130 μL droplet of the PDMS mixture was placed onto each of the plasma-treated glass coverslips. The PDMS droplets were allowed to spread across the glass coverslips on a level countertop for at least 30 minutes, resulting in a PDMS layer that is approximately 250 μm in height. The PDMS-coated coverslips were degassed for 30 minutes before transferring to a 65 °C oven overnight to cure. The following day, the PDMS substrates were removed from the oven and cooled at room temperature. To extract unpolymerized monomers, the PDMS substrates were submerged in 100% ethanol for at least 3 hours, followed by multiple rinses with DI water before drying in a 65 °C oven overnight.

### 2.4. Microcontact printing and functionalization of black dots

The patterned PDMS stamps and PVA film were used to deposit a layer of fluorescent protein onto the flexible PDMS substrates similar to previously published techniques[29,30] (Fig. 1B). All steps were performed at room temperature and preferably in a standard tissue culture hood. First, Alexa-Fluor 488, 594, or 647-conjugated-BSA (5 mg/mL, Life Technologies) was diluted 1:2000 in PBS (1X without calcium or magnesium, Life Technologies), and a 400 μL droplet was gently placed onto a patterned stamp (about 1 cm^2^ area) within a petri dish. The droplet was left on the stamp for 30 minutes to allow the fluorescent BSA to adsorb onto the surface. Fresh PBS was slowly added to the petri dish until the liquid level rose above the stamp. The stamp was removed from the PBS and rinsed 3 times in fresh PBS dishes by gently submerging the stamp. After the final rinse, the stamp was dried with a gentle stream of nitrogen gas.

Next, the PVA film was used to transfer the fluorescent pattern from the stamp to the flexible substrate. A PVA film was trimmed to a size slightly larger than the stamp. The film was plasma treated for 60 seconds to facilitate protein transfer from the stamp. Using a pair of tweezers, the film was then lowered onto the dried stamp. The film was gently pressed onto the stamp using rounded-tip tweezers to remove any air gaps, and a thin piece of glass slide was placed on top of the film. A 50-gram weight was placed onto the glass slide to maintain close contact between the film and stamp. After 20 minutes, the weight and glass slide were removed and the PVA film was gently peeled from the stamp and transferred to the flexible PDMS substrate. Again, rounded-tip tweezers were used to gently press the film onto the substrate and remove any air gaps. The film was left on the flexible PDMS substrate for 20 minutes. The substrate was then submerged in PBS for up to 5 minutes, causing the film to rehydrate and float away from the surface where it can be discarded. The final substrate containing the pattern of fluorescent BSA, dubbed “black dots,” was stored in PBS overnight at 4 °C before cell seeding and can be stored for at least 1 week.

On the day of cell seeding, von Willebrand Factor (VWF) (Haematological Technologies) was diluted in PBS to 5 μg/mL and was pipetted onto the black dots. To encourage droplet spreading over the black dot surface, a glass coverslip was gently placed on top of the droplet. The von Willebrand Factor was incubated for 1-1.5 hours at room temperature before the coverslip was removed. To block the surface, the substrate was then submerged in a 0.2% Pluronic F-127 (BASF) in PBS for 30 minutes. The substrate was finally submerged into PBS and stored until platelet seeding.

To quantify VWF adsorption onto the surface, black dots with and without VWF treatment were blocked with 10% goat serum (Life Technologies, diluted in PBS) for 1 hour and then incubated with a FITC-labeled anti-von Willebrand Factor antibody (Abcam, ab8822) for 1 hour). Substrates were mounted onto glass coverslips using Fluoromount-G mounting medium (Life Technologies) for confocal microscopy. Images collected with the same settings were quantified using MATLAB to characterize FITC fluorescence (and therefore VWF adsorption) on the fluorescent BSA and the non-fluorescent black dots for substrates with and without VWF treatment.

### 2.5. Platelet isolation and seeding

Platelet-rich plasma (PRP) was collected from consenting research participants by plateletpheresis using the Trima Accel^®^ automated collection system. Research participants were healthy and not taking any platelet inhibiting medications. Platelets were isolated from plasma by platelet centrifugation washing modified from previously described protocols[20]. Platelets were pelleted at 1000 g and resuspended in HEN Buffer, pH 6.5 containing 10 mM HEPES (Sigma), 1 mM EDTA (Corning), and 150 mM NaCl (Fisher Scientific) and supplemented with 0.5 μM prostacyclin (PGI_2_) (Sigma). To prevent activation, platelets were incubated for 10 minutes at room-temperature and then repeat treated with 0.5 μM PGI_2_ and pelleted via centrifugation at 800 g. Platelets were resuspended and diluted to 3∙10^8^ platelets/mL with modified Tyrode’s buffer, pH 7.3 containing 5 mM HEPES (Sigma), 137 mM NaCl (Fisher Scientific), 5.5 mM glucose (Fisher Scientific), 12 mM NaHCO_3_ (Sigma), 0.3 mM NaH_2_PO_4_ (Sigma), 2 mM KCl (JT Baker), 1 mM MgCl_2_ (Sigma), and 2 mM CaCl_2_ (Macron Fine Chemicals) and supplemented with 0.35% (w/v) human serum albumin and 0.02 U/mL apyrase.

Immediately before seeding the washed, isolated platelets onto the black dots, the platelets were further diluted to 2.5∙10^7^/mL in Tyrode’s Buffer, pH 7.5 containing 10 mM HEPES (Fisher Scientific), 138 mM NaCl (JT Baker), 5.5 mM glucose (ACROS Organics), 12 mM NaHCO_3_ (Sigma), 0.36 mM NaH_2_PO_4_ (Sigma), 2.9 mM KCl (VWR), 0.4 mM MgCl_2_ (Fisher Scientific), and 0.8 mM CaCl_2_ (VWR International). After dilution, 10 million platelets were seeded onto each black dot substrate. To allow time for initial platelet binding onto the black dots, platelets were incubated at room-temperature for 10 minutes. To remove unattached platelets, black dots were then gently dipped in PBS and then immediately submerged in fresh Tyrode Buffer. To allow time for platelet adhesion and contraction on the black dots, the platelets were incubated for an additional 30 minutes at room-temperature. Incubation times were selected to reduce temporal differences in platelet contraction by 1) preventing new platelet binding after 10 minutes, such that all platelets were on the surface for 30-40 minutes and 2) allowing platelet binding and contraction for 30 minutes so that platelets reach a maximum contraction[19,21,25].

### 2.6. Immunocytochemistry

Platelets were fixed with 4% paraformaldehyde for 20 minutes, permeabilized with 0.1% Triton X-100 for 10 minutes at room temperature, and blocked with 10% goat serum (Life Technologies, diluted in PBS) for 1 hour. Platelet F-actin was labeled with phalloidin 488 (Life Technologies), and platelet GPIb was labeled with a CD42b monoclonal antibody, clone SZ2 (Life Technologies) and a goat anti-mouse IgG secondary antibody (Life Technologies). Substrates were mounted onto glass coverslips using Fluoromount-G mounting medium (Life Technologies) for confocal microscopy.

### 2.7. Imaging and image analysis

Fixed and stained platelets were imaged on a Nikon A1R or a Leica SP8 confocal microscope with a 60x oil objective (NA = 1.4). Images of platelets were taken with a large enough field of view to ensure several black dots with no deformation surrounded each platelet.

To quantify the deformation of the black dots, we modified a previously existing method for tracking objects[31]. The fluorescent image of the black dots was first run through a spatial bandpass filter with a characteristic noise length scale of 1 pixel. The dots were identified with a peak finding algorithm using a peak threshold value of 0.15. For each dot, the centroid was then found to subpixel accuracy. To calculate the displacement of each dot, the zero-displacement state of the black dots must be determined. The dots in the image were organized into a rectangular array of rows and columns. For each row and column, a line was fit through four of the dots near the edges of the image. We assume that the dots near the edge of the image have little or no displacement and can therefore be used as reference for the highly displaced dots near the cell. The lines fit through the rows intersect with lines fit through the columns, and the intersection points were used as the zero-displacement state. From here, the displacement of each dot was calculated by subtracting the zero-displacement position from the deformed position.

### 2.8. Force calculation

Traction forces are calculated from the surface displacements using regularized Fourier Transform Traction Cytometry (FTTC)[32,33]. FTTC requires a rectangular grid of displacements which is obtained trivially by our black dots pattern without any interpolation. Any missing data locations near the periphery or corners of the image are filled in with imaginary data points assigned with zero displacement. A regularization parameter of 5∙10^−8^ was used to smooth out noise in the traction forces.

To assess whole-cell contractility, we calculate the total force and net force for each cell. Total force is calculated by summing the force magnitudes from each dot underneath the cell. Net force is similarly calculated by simply adding the force vectors from each dot together; the cell is assumed to be in static equilibrium, so the net force should tend towards 0. Because the cell can adhere to spaces between black dots, we consider all black dots within the cell boundary and within 1 dot spacing outside the cell boundary for these calculations.

### 2.9. Area and circularity calculations

The cell boundary and area were determined in MATLAB using a user-adjusted threshold and shape fill on the fluorescent F-actin image. We also tested determining the cell boundary using the GPIb fluorescent image and no difference in cell boundary was observed. Circularity was calculated from the cell boundary using: *C* = 4π*A*/*P*^2^, where *C* is the circularity, *A* is the area, and *P* is the perimeter[34]. Circularity values can range from 0 to 1, where a value of 1 indicates a perfect circle.

### 2.10. F-actin dispersion calculation

For each cell, the stained F-actin image was normalized such that the fluorescent intensity within the cell boundary spanned values 0 to 1. A threshold was set at 0.1, and the F-actin dispersion was calculated as the percentage of cell pixels above this threshold. Based on this calculation, a cell with well-dispersed or uniform F-actin stain will receive a value closer 1, while a cell with localized F-actin intensity will receive a value closer to 0. The threshold value of 0.1 was chosen because it resulted in both the largest spread of F-actin dispersion between cells and yielded a quantification most consistent with qualitative measures.

### 2.11. K-means clustering

A K-means clustering analysis was utilized to separate the data into clusters in an unbiased way. First, the area, circularity, and F-actin dispersion data were normalized. The built-in MATLAB function “kmeans” was used to cluster the data based on the area, circularity, and F-actin dispersion. Cell force was not included for purposes of clustering. To determine the optimal number of clusters, the built-in MATLAB function “evalclusters” was employed for up to 6 clusters using either Silhouette or Gap evaluation criteria with default settings. The optimal number of clusters is defined as the one with the highest Silhouette value or as the lowest number of clusters such that the mean Gap value for the next highest number of clusters falls within the standard error of the previous one. Silhouette and Gap values were evaluated for up to 6 clusters, and both criteria suggest that 2 clusters is optimal for our data set (Supplementary Fig. 10).

### 2.12. Statistics

To compare donor to donor variability, a one-way ANOVA and Tukey’s post hoc test was used to determine whether differences in the means between donors were statistically significantly significant.

To examine effects of area, circularity, and F-actin dispersion on force, each covariate was centered at its mean and circularity and F-actin dispersion were transformed to a 0-100 scale. This transformation does not affect the results. A multivariate mixed effects model with random donor effects was used to analyze the centered data and determine the influence of individual covariates and interactions between covariates.

To determine if forces of platelets in cluster 1 and cluster 2 (as determined by the K-means clustering analysis) were significantly different from each other, a student’s t-test was used. For all tests, significance is considered p < 0.05.

### 2.13. Cell exclusion considerations

To reduce systematic error in our data, we have several considerations for excluding cells from the analysis. Our analysis requires all black dots near the edge of the cell field of view to be undeformed; any cells within close proximity of each other will disrupt this requirement. Therefore, cells which are close in proximity to other cells are automatically disregarded from all analysis; close proximity is defined here as two neighboring cell boundaries coming within 2 μm of each other. Another exclusion criterion are high net forces which are indicative of irregular patterning or mounting issues. We chose a cutoff of 10 nN and excluded all cells exhibiting net forces higher than this cutoff from our data sets. Additionally, we excluded platelets that did not spread by excluding platelets < 10 μm^2^ that had no filopodial or lamellipodial protrusions. Finally, fluorescence from F-actin staining was observed to be highly variable in some cells; we tuned the exposure time to the best of our abilities but for some cells it was difficult to completely eliminate image saturation. Therefore, we excluded cells that had greater than 1% saturated pixels within the cell boundary from the analysis shown in Figures 3, 4, and 5.

## 3. RESULTS

### 3.1. Microcontact printing of black dots with uniformity in size, spacing, and shape

Black dots were manufactured, coated with extracellular matrix protein (ECM), and seeded with platelets (Fig. 1B). In this work, the VWF was chosen as the ECM to facilitate platelet adhesion. Using a fluorescent anti-VWF antibody, we visualized VWF adsorption on the surface and found that VWF is adsorbed contiguously across the surface, with some preference for binding to the fluorescent BSA (Supplementary Fig. 1A-G). Additionally, we have coated the black dots with other ECM such as fibrinogen (Supplementary Fig. 1H-J) and laminin (Supplementary Fig. 1K-M), so many cell types can be studied with this technique.

Once the black dots technique is optimized, it provides a consistent pattern and can be tailored to suit the nature of the cells. We created black dots with BSA conjugated with Alexa Fluor 488, 594, and 647 to demonstrate the versatility in the fluorescent coatings that are possible (Fig. 1C). Through quantitative image analysis, the black dots were found to be uniform in size (1.02 ± 0.03 μm diameter), spacing (1.96 ± 0.02 μm center-to-center), and shape (0.93 ± 0.01 circularity) (Fig. 1D). This pattern uniformity is critical for obtaining accurate results from image analysis and force calculations.

The soft PDMS we used resulted in a substrate with a stiffness of 13.5 kPa and was selected because it was physiologically relevant for platelets[35,36]. We tested softer and stiffer mixtures of Sylgard 527 and 184, but they were not optimal for traction force measurements with platelets because the resulting deformations of a subset of platelets were too large or too small to measure accurately (Supplementary Fig. 2). For measurements with other cell types, the ratio of PDMS mixtures can be adjusted to match their level of contractility.

We have found that microcontact printing for the black dots can be a sensitive process, so care must be taken in preparing and storing the substrates. We have provided helpful tips and avoidable pitfalls for others to refer to in adopting the technique (Supplementary Note). Using our protocol, we typically print areas of black dots of 1 cm^2^, but we have printed areas up to nearly 10 cm^2^ with a larger PDMS stamp (Supplementary Fig. 3). The black dots approach could potentially be scaled to larger culture dishes for even higher throughput in measurements. Overall, we have shown that the microcontact printing and sacrificial film technique can deposit fluorescent BSA patterns of black dots with regular size, spacing, and shape that cover a large surface area for experiments with cells.

### 3.2. Reference-free Traction Force Microscopy with black dots

To demonstrate the black dots approach, we seeded human platelets and measured their traction forces. Washed platelets were seeded onto VWF-coated black dots for 10 minutes to allow them to adhere and then rinsed gently to remove unbound platelets. We waited an additional 30 minutes to allow the platelets to spread and contract before fixing the samples. This timing for platelet binding and contraction was selected based on dynamics of platelet force generation[12,19,21,24]. With immunofluorescence staining and confocal microscopy, many platelets can be captured in a single image (Fig. 2A and Supplementary Fig. 4). We note that the platelets had various shapes and sizes similar to previous observations on glass substrates[24,37]. Platelets were also seeded onto black dots without VWF and we observed that platelets did not bind and spread, demonstrating that the platelet adhesion is VWF-specific.

**Figure 2.**
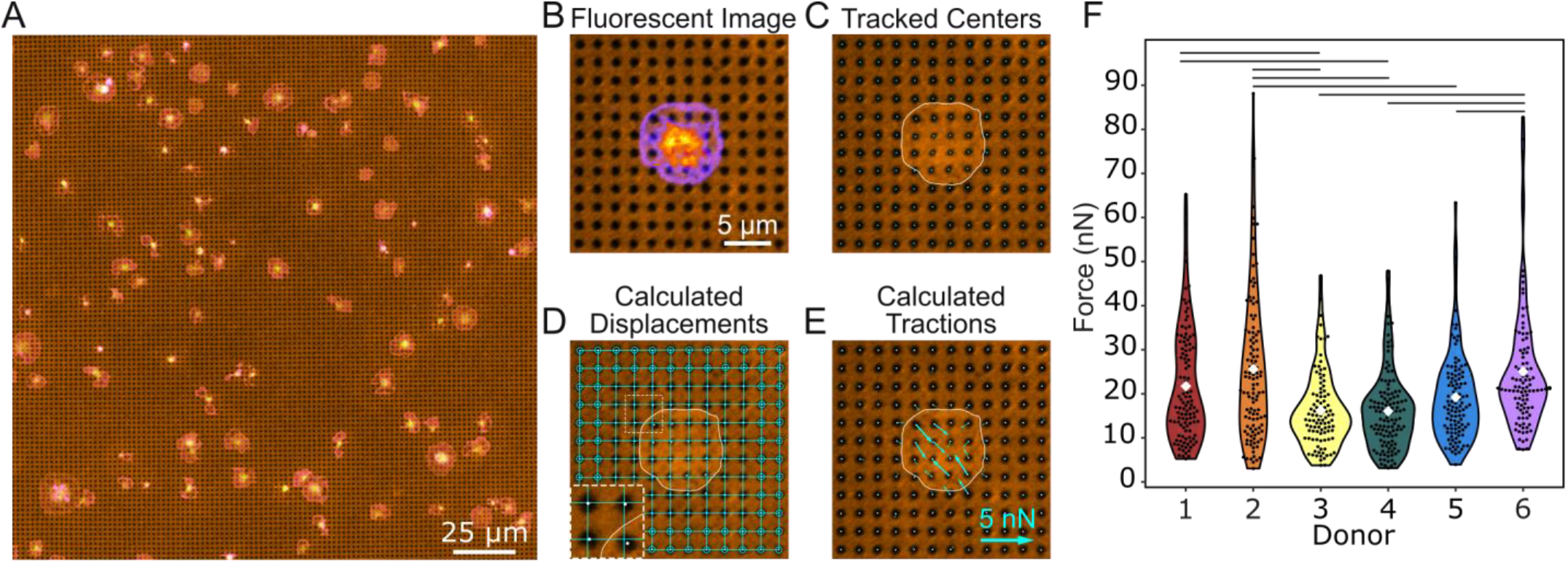
Black dots offer a higher yield way to measure forces. (A) Example of a field of view containing many platelets adhered to and contracting on the black dots. Here, platelets are stained for both F-actin (green) and GPIb (magenta). (B) A fluorescence image of deformed substrate with platelet stained for F-actin (pixel intensity scaled from purple to yellow). (C) The black dots pattern is binarized and the centroid of each dot is found using automated detection. (D) Undeformed dots near the edges (circled) are used to fit horizontal and vertical lines throughout the entire field of view. The intersection of these lines marks the zero-displacement state of each dot. Inset is 2x magnified. (E) Forces are calculated from the displacement of each dot relative to its zero-displacement (undeformed) location. (F) Forces from at least 100 platelets from 6 donors show high variability within each donor and between donors. Lines indicate significant differences in donor forces (p < 0.05 when tested with a one-way ANOVA and Tukey’s post hoc test). Number of cells analyzed for donors 1, 2, 3, 4, 5, and 6 are n = 111, 117, 100, 120, 112, and 100, respectively.

Individual platelets within an image are cropped and analyzed separately (Fig. 2B). The centroid of each black dot is identified using automated detection (Fig. 2C). Black dots at the edge of the region are used to form a grid of best-fit lines whose intersections denote the undeformed position of each black dot (Fig. 2D). For each black dot, the distance from its centroid to its respective intersection in the zero-displacement grid (Fig. 2D inset) is used to calculate the magnitude and direction of the forces using regularized Fourier Transform Traction Cytometry (FTTC) (Fig. 2E)[32,33]. The black dot technique is suited well for FTTC, which requires the measured displacements to be on a regular grid. The total force for each platelet is calculated by summing the force magnitudes of each dot under the cell. All data plotted in this work is the total force of single platelets.

The total contractile forces of platelets from six healthy donors were analyzed (Fig. 2F). The mean force measured by black dots was 24.1 nN, which is similar to other methods that have reported forces between 19 and 200 nN for individual platelets[12,21,25,26]. We also noted a wide range of forces, from 3.5 to 98.7 nN (28-fold difference) with a standard deviation of 14.7 nN. This observation agrees with other studies; atomic force microscopy found that platelet forces varied from 1.5 to 79 nN[25] (more than 50-fold difference), and subsequent work using TFM and nanoposts have also observed heterogeneity in platelet forces[21,26]. We observed heterogeneity both within and between donors, including a standard deviation of 13.7 nN among platelets from the same donor as well as statistically significant differences between mean platelet force from different donors (lines in Fig. 2F). Our results show that platelet forces measured with black dots are similar in magnitude to previous measurements and indicate that populations of platelets produce a wide range of forces.

### 3.3. Platelet forces correlate with spread area, circularity, and F-actin dispersion

We questioned whether the heterogeneity in total platelet forces could be attributed to their size, shape, and/or cytoskeletal structure. Platelets typically bind to a surface, increase their spread area approximately 5-fold over about 10 minutes, and then sustain their maximum spread area[21,23,24]. In our experiments, platelets were allowed to adhere and spread for 30-40 minutes after attachment to allow them to reach their maximum area. We examined spread area as a factor influencing the overall magnitude of traction forces in platelets as it has been observed previously[21,22] as well as in many other cell types[38–40]. We find that the spread area of platelets ranged from 8.7 to 205.5 μm^2^, with a mean and standard deviation of 43.5 ± 22.4 μm^2^. We observed a positive relationship between force and area, having a best-fit slope of 0.53 nN/μm^2^ (R^2^ = 0.49) (Fig. 3A-C). This force-area relationship is maintained in all six donors, with some minor differences between them (Supplementary Fig. 5). While our results indicate a strong force-area relationship in platelets, we do find a degree of heterogeneity in our results. For example, platelets with a spread area of 50-55 μm^2^ exerted forces from 14.3 to 71.0 nN, with a mean and standard deviation of 31.8 ± 13.3 nN. Although spread area has a strong correlation with platelet forces, it does not fully account for their contractile output.

**Figure 3.**
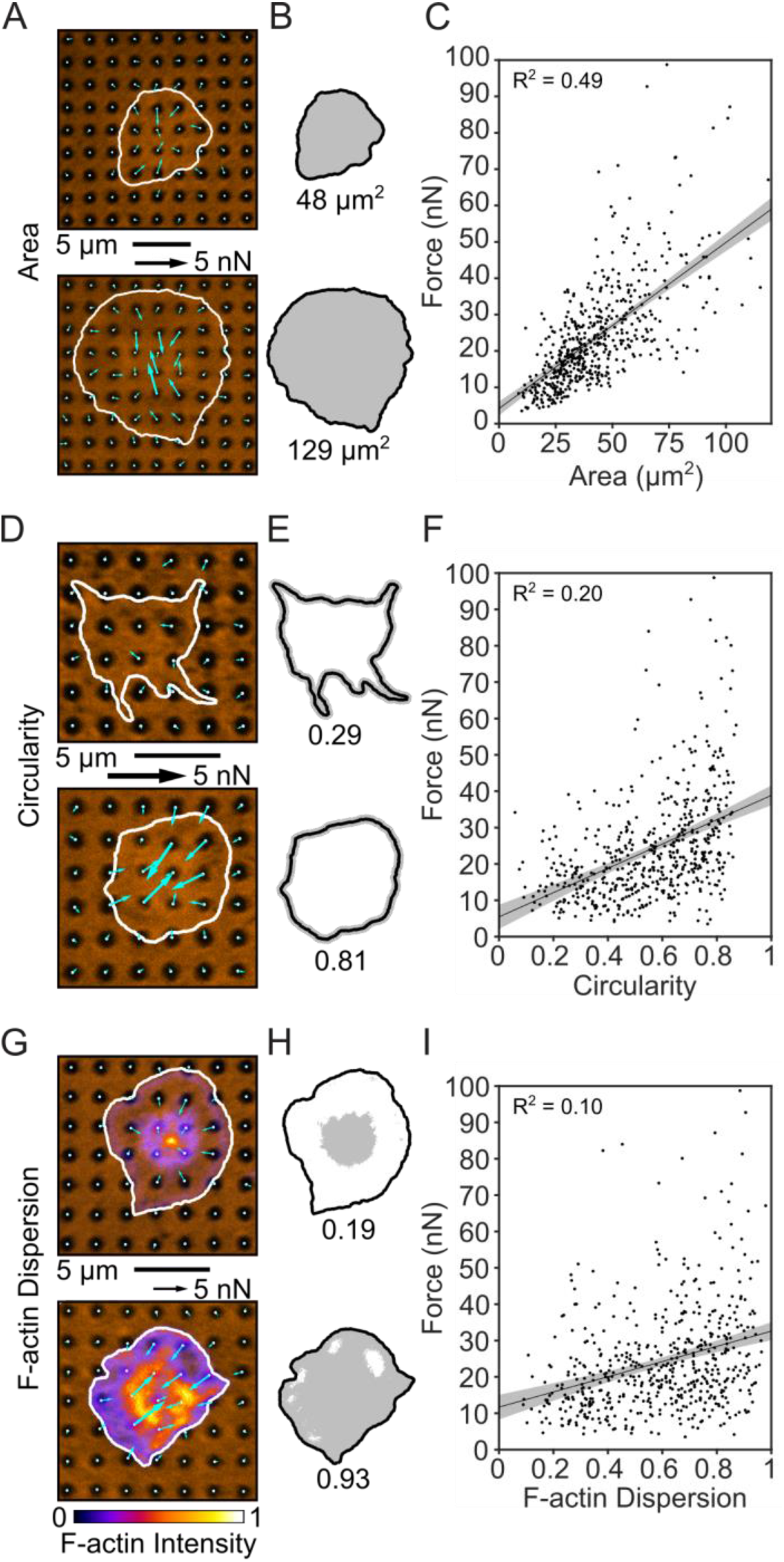
Platelet size, shape, and structure correlate with force. (A) Two examples of platelets with small and large areas. (B) Cell boundary and area measured from (A). (C) Platelet forces and area are linearly related. Note that in this panel, the x-axis maximum is zoomed to better view the data. Due to this axis zoom, two points (0.37% of the data) are not shown, but all points are included within all analyses (including the fit line calculation). (D) Two examples of platelets with low and high circularity. (E) Cell boundary and circularity measured from (D). (F) Platelet force and circularity show a moderate positive relationship. (G) Two examples of platelets with low and high F-actin dispersion. Color bar indicates fluorescence intensity which has been normalized to calculate F-actin dispersion. (H) Cell boundary and F-actin dispersion measured from (G). (I) F-actin dispersion is moderately positively correlated with platelet force. Shaded regions of fit lines indicate 95% prediction interval for the data.

Another aspect we considered was the dramatic shape changes of platelets such as their transition from discoid to spherical shape upon activation and their extension of filopodial protrusions in the early stages of platelet spreading[23,41]. Because these shape changes are important to platelet function, we investigated whether platelet shape correlates with force. We observed that adherent platelets on the black dots adopt a variety of different shapes, ranging from stellate to circular. We used an image analysis metric of circularity to quantitatively assess these different shapes (Fig. 3D-E). We find that more circular platelets generate larger forces than ones that are stellate and less circular ones (Fig. 3F). However, the best-fit slope of this relationship is 0.33 nN/0.01 circularity units (R^2^ = 0.20) so it is not as correlative as the force-area relationship. All six donors showed similar force-circularity behavior (Supplementary Fig. 6). These results indicate that circularity has a moderate correlation with force.

Due to the underlying role of actin remodeling in initiating platelet shape changes and generating cellular forces[23,41], we used black dots to determine whether actin arrangement correlates with platelet forces. When we stained the platelets to view their F-actin network on black dots, we observed a cortical actin ring around the cell boundary of most platelets. However, there were some distinct differences in the F-actin structure in their interior, ranging from punctate to dispersed (Fig. 3G-H). The amount of F-actin dispersion was quantified and plotted against force. We found that platelets with more dispersed F-actin structure typically generated higher forces and the best-fit slope of this relationship is 0.21 nN/0.01 F-actin dispersion units (R^2^ = 0.10) (Fig. 3I). All six donors exhibited a similar force-F-actin dispersion relationship (Supplementary Fig. 7). Collectively, we find that area has a strong correlation with force, and that circularity and F-actin dispersion moderately correlate with force.

### 3.4. Multivariate mixed effects modeling reveals cooperative effects between F-actin dispersion and circularity and between F-actin dispersion and spread area

Because platelet size, shape, and cytoskeletal organization change concurrently during platelet adhesion[23], we also investigated whether area, circularity, F-actin dispersion correlate with each other (Fig. 4A, D, H). For area and circularity, we observed a moderate correlation (R^2^ = 0.26) (Fig. 4A). To visualize the combined relationship of circularity and area with increasing force, platelets were split into four equally sized groups by force, i.e., quartiles. Notably, low-force platelets had small areas, but also had a wide range of circularities. On the other hand, high-force platelets almost exclusively had high area and high circularity (Fig. 4B and Supplementary Fig. 8 to see graphs with all points). By plotting the median of each quartile, we observed that circularity and area increase together with increasing force (Fig. 4C). Similarly, area and F-actin dispersion (Fig. 4D-F) as well as F-actin dispersion and circularity (Fig. 4G-I) increase together with each force quartile. The particularly extreme shift observed in the circularity versus F-actin dispersion contour plots is somewhat surprising, given that each of these factors only moderately correlate with force. These results indicate that there are some correlations between platelet area, circularity, and F-actin dispersion and that together, they have strong effects on force. We next turned to a more robust approach to assess interaction effects between these factors.

**Figure 4.**
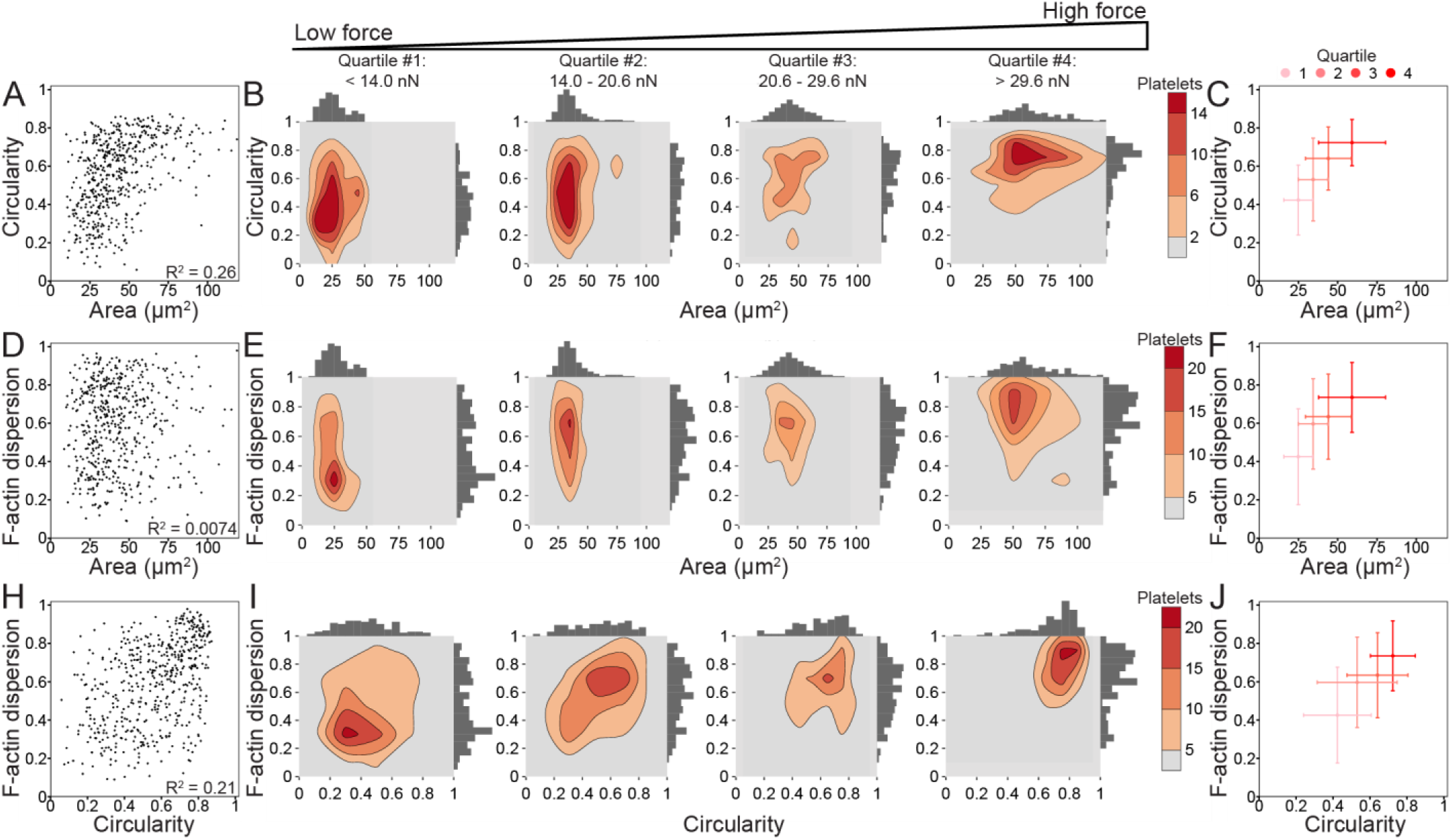
Platelet size, shape, and structure do not strongly correlate with each other, but increase together with force. (A) Area and circularity moderately correlate (R^2^ = 0.26) when plotting platelets of all forces (n = 540). (B) To examine the relationship of circularity and area with increasing force, platelets are split into four quartiles (n = 135 in each quartile) by force, where the lowest force quartile (quartile #1) includes platelets generating less than 14.0 nN force, quartile #2 contains platelets generating 14.0-20.6 nN force, quartile #3 contains platelets generating 20.6-29.6 nN force, and the highest force quartile (quartile #4) contains platelets generating greater than 29.6 nN force. Contour density plots and histograms of area and circularity at each force quartile show that quartile #1 (low-force platelets) have low area and a large range of circularity, while quartile #4 (high-force platelets) tend to have both higher area and high circularity. (C) The median and median standard deviation show that circularity and area increase together with each force quartile. (D) F-actin dispersion and area do not correlate (R^2^ = 0.0074), (E-F) but show a similar trend when examining force quartiles. (H) F-actin dispersion and circularity moderately correlate (R^2^ = 0.21) and (I-J) show a shift from low-force platelets having large ranges of circularity and F-actin dispersion to high-force platelets have high circularity and high F-actin dispersion. Note that in A-F, the x-axis maximum is zoomed to better view the data. Due to this axis zoom, two points (0.37% of the data) are not shown, but all points are included within all analyses.

To further investigate how F-actin dispersion, circularity, and area affect force in different but overlapping ways, we used R Studio to create a multivariate mixed effects model allowing for 2-way interactions between each of the factors (Table 1). The mixed effects model shows that across donors, the difference in force between two platelets that differ in area by 1 μm^2^ (while other factors remain constant) is 0.41 nN (95% CI: 0.37, 0.45) on average, with the larger platelet generating more force (Table 1). Similarly, when holding other factors constant, two platelets that differ in circularity or F-actin dispersion by 0.01 will respectively differ in force by 0.069 nN (95% CI: 0.02, 0.11) and 0.15 nN (95% CI: 0.11, 0.19), on average. From estimates and standard errors in Table 1, p-values are calculated to determine what factors and interactions have a significant (p < 0.05) effect on force. All individual factors (area, circularity, and F-actin dispersion) significantly contribute to force when controlling for the other factors. In addition to these main effects, two interaction terms were significant at the p < 0.05 level: F-actin dispersion interacting with area and F-actin dispersion interacting with circularity, each of which is a positive, cooperative effect. For example, when circularity is average, F-actin dispersion and force have a positive relationship with a slope of 0.069 nN/0.01 F-actin dispersion units. When circularity is one standard deviation above average, the relationship between F-actin dispersion and circularity is stronger and has a slope of 0.12 nN/0.01 F-actin dispersion units (75% increase). Conversely, when circularity is one standard deviation below average, the relationship between F-actin dispersion and circularity is weaker and has a slope of 0.017 nN/0.01 F-actin dispersion units (75% decrease) (Supplementary Fig. 9E and Supplementary Fig. 9 for all other interaction plots). Area interacting with circularity has a p-value of 0.2079 and is not significant at the p < 0.05 level. This multivariate mixed effects model supports the contribution of area, circularity, and F-actin dispersion to force and suggests a complex relationship between these factors. Additionally, this analysis reveals significant cooperative effects between F-actin dispersion and circularity and between F-actin dispersion and area.

**Table 1.** Multivariate mixed effects model summary shows significant interaction effects. The estimate column indicates the expected increase in force (in nN) if the variable in that row increases by 1 μm^2^ (area) and/or 0.01 units of circularity or F-actin dispersion while all other factors remain constant. The standard error column indicates the error of that estimate. From these estimates and standard errors, p-values are calculated to determine what factors and interactions have a significant (p < 0.05) effect on force. In addition to these individual effects, pairs of effects were tested for interactions, where a positive value in the estimate column indicates a cooperative effect and a negative value an antagonistic effect.

|  | **Estimate** | **Std. Error** | **p-value** |
| --- | --- | --- | --- |
| **Area** [nN/$\mu\text{m}^2$] | 0.4100 | 0.0209 | < 0.001 |
| **Circularity** [nN/0.01 units] | 0.0687 | 0.0247 | 0.0053 |
| **F-actin dispersion** [nN/0.01 units] | 0.1488 | 0.0187 | < 0.001 |
| **Circularity : F-actin dispersion** [nN/(0.01 units*0.01 units)] | 0.0024 | 0.0010 | 0.0134 |
| **Area : F-actin dispersion** [nN/($\mu\text{m}^2$*0.01 units)] | 0.0024 | 0.0008 | 0.0028 |
| **Area : Circularity** [nN/($\mu\text{m}^2$*0.01 units)] | -0.0012 | 0.0009 | 0.2079 |

### 3.5. Unbiased clustering supports relationship between spread area, circularity, F-actin dispersion, and platelet force

Big data analyses are powerful tools that can help extract significant information in data sets that are large and unwieldy. In our population of platelets, we observed a large range of shapes, sizes, and structures, so we wanted to investigate whether there are clusters or subpopulations of platelets in our data set. We performed an unbiased K-means clustering analysis on platelet area, circularity, and F-actin dispersion to locate possible clusters, and to see if the relationships we observed between these factors and force could be explained by distinct clusters or subpopulations of platelets. Two clusters arose from this analysis: cluster 1 is generally characterized by low spread area, circularity, and F-actin dispersion while cluster 2 is high spread area, circularity, and F-actin dispersion (Fig. 5A-D and Supplementary Fig. 10A-D). We also performed the clustering analysis on each donor individually; each donor generally formed two clusters that were similar to the two clusters in the overall data set (Supplementary Fig. 10E-J). For this clustering analysis, we intentionally did not include the force data; despite this agnostic approach to platelet force, we find that cluster 2 has significantly higher forces than cluster 1 using a student’s t-test (Fig. 5E), supporting our earlier findings.

**Figure 5.**
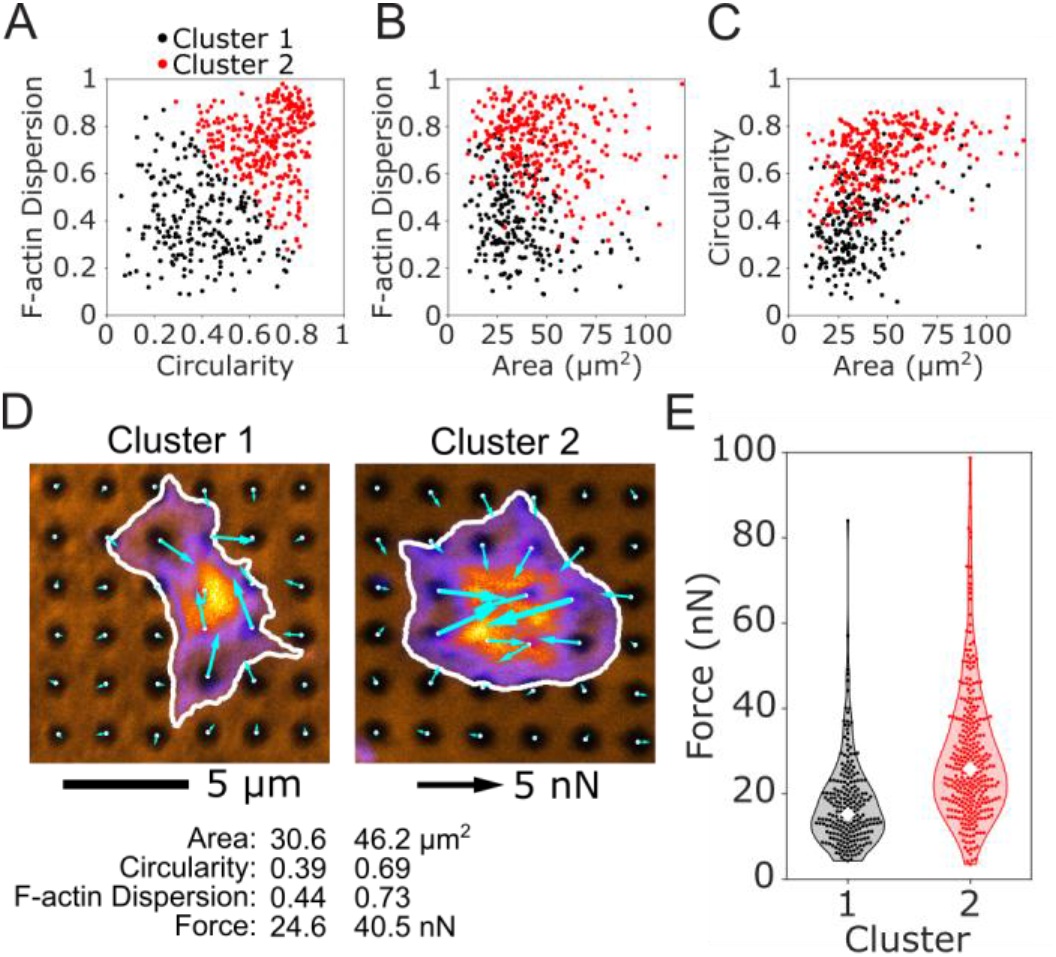
K-means clustering of platelet size, shape, and structure predict differences in force. An unbiased K-means clustering approach based on platelet area, circularity, and F-actin dispersion separated the population of platelets into 2 clusters. The two clusters are shown for (A) F-actin dispersion and circularity, (B) F-actin dispersion and area, and (C) circularity and area. (D) The most representative platelet from each cluster is shown. Platelets from cluster 1 have smaller area, lower circularity, and lower F-actin dispersion than cluster 2. (E) Forces from cluster 2 are significantly higher than cluster 1, even though force was not used to determine the clusters.

K-means clustering was chosen here due to its simplicity and widespread use, although other clustering methods may be more appropriate depending on the data set. By eye, the two clusters in our data set tend to lie on a continuum rather than distinct clusters with no overlap. This could indicate that the two clusters do not originate from distinct sources, but instead emerge from a single gradient such as platelet age in circulation, where older platelets tend to have less spread area, less circularity, and less dispersed F-actin. The physiological origin and significance of these clusters will be further investigated in future studies. Ultimately, this clustering analysis serves as a demonstration of big data analyses that are made possible by data from hundreds of cells collected with a high-yield method.

## 4. DISCUSSION

Here, we showed how the black dots approach is used to measure traction forces in platelets. The black dots were coated with VWF, an adhesive blood protein that mediates platelet adhesion. The choice of ECM depends on the cell type being studied, so we demonstrated that black dots could instead be coated with fibrinogen or laminin which are commonly used for many cell types. Additionally, the substrate stiffness of 13.5 kPa was selected because it was physiologically relevant for platelets[35,36]. Other cell types may be more suited to a different stiffness; our platform utilizes a mixture of two types of PDMS which can be adjusted to change the final substrate stiffness[29]. Overall, the black dots approach may be useful to measure traction forces for many cell types, and not only platelets.

We used the black dots technique to characterize the relationship of force with platelet size, shape, and cytoskeletal structure. We measured forces of more than 500 platelets, which is five times more than previous studies[19,21,22] and is on par with existing high-yield methods that directly control cell shape and area[12]. The magnitude of forces from our technique is similar to other methods that have reported forces for individual platelets[12,21,25,26]. For the first time, we were able to correlate platelet forces to platelet circularity and F-actin dispersion. This was only possible with the black dots technique because it does not constrain platelet shape or size and is compatible with the immunofluorescent techniques necessary to study the cytoskeleton. We found significant associations between spread area, circularity, F-actin dispersion, and force, as well as interactions between these factors that significantly contribute to platelet force generation. When the independent effects are determined with a multivariate mixed effects model, F-actin dispersion associates more strongly with force than circularity, because it is less correlated with area. Moreover, cooperative interactions between F-actin dispersion and both area and circularity further highlight the importance of F-actin structure in generating contractile forces and provide new insight into the large heterogeneity of observed platelet forces.

Beyond the measures of area, circularity, and F-actin dispersion, the amount of contractile force a cell can generate likely depends on several factors that we have not measured here, including activation of the actomyosin network by phosphorylation, amount and organization of the contractile fibers, genetic differences between donors, and disease states. We anticipate that the black dots platform may be used in conjunction with detailed fluorescent staining, western-blotting, or genetic screening to further enhance the understanding of force generation of platelets and other cells.

## 5. CONCLUSION

The black dots approach is a high-yield single-cell force measurement platform that is compatible with fixed cells without constraining cell shape and size. It relies on microcontact-printing and algorithms from reference-free traction force microscopy to measure traction forces of individual cells. We demonstrate the technique’s benefits by measuring forces of more than 500 platelets, a high yield for traction force measurements. Using this approach, we were able to correlate platelet forces to platelet circularity and F-actin dispersion, revealing cooperative effects. By tuning the substrate stiffness, extracellular matrix protein, and BSA fluorescence, the black dots approach may be useful to measure the forces in many cell types beyond platelets.

## ACKNOWLEDGEMENTS

This work was supported by the National Science Foundation (CMMI-1661730, CMMI-1824792), the National Institutes of Health (EB001650, HL147462, HL149734, GM135806, AR074990, TR003519, DE029827), and the Institute for Stem Cell and Regenerative Medicine Fellows Program. Imaging in this study was completed in the Lynn & Mike Garvey Imaging Core with the helpful guidance of Dale Hailey. The Department of Biostatistics Statistical Consulting Services and Prof. Megan Othus assisted with the statistical analysis for this study. We would also like to thank Robin Zhexuan Yan, Kenia Diaz, Francisco Morales, and Anabela Soto for their assistance testing the robustness of black dot manufacturing and/or the usability of the black dot analysis code.

## AUTHOR CONTRIBUTIONS

K.M.B. and M.Y.M. contributed equally to this study. K.M.B., A.L., and N.J.S. conceived of the black dot method. K.M.B., M.Y.M., and A.L. optimized the black dot method. J.M. recruited blood donors, collected, and washed platelets. M.Y.M. performed the platelet force assay. K.M.B., S.J.H., A.H., and N.J.S. determined the model to calculate force from black dot displacement. K.M.B. primarily wrote the analysis code, with contributions from M.Y.M. and Z.S. M.Y.M. and K.M.B. analyzed the images, plotted the data, and made the figures. J.H., M.Y.M., and K.M.B. conducted the statistical analyses. K.M.B, M.Y.M., W.E.T., and N.J.S. designed the experiments, interpreted the data, and wrote the manuscript. All authors reviewed and edited the manuscript.

