## Supplemental text and figures for "Black Dots: Microcontact-Printed, Reference-Free Traction Force Microscopy"

**Supplementary Material**

### 1. Notes

Throughout the development of the black dots method, we encountered steps several steps in the manufacturing process that required optimization. Here are some tips that increased the reliability when manufacturing black dots:

Master mold

- For platelets, the master contained a designed pattern of round dots with a nominal diameter of 1 µm and spacing of 2 µm. The actual spacing between dots on the SU-8 master mold was found to be 1.99 µm measured by widefield microscopy (data not shown here). The final fluorescent pattern of black dots were measured to have a mean diameter of 1.02 µm and mean spacing of 1.96 µm as shown in Figure 1D, suggesting that the negative PDMS stamp had some shrinkage during the manufacturing steps.

Soft PDMS substrates

- When making the soft PDMS coverslips, keeping the platform flat to ensure even spreading of PDMS was important to reduce tilt when imaging. We found that placing the coverslips onto the inside of the top lid of a petri dish, rather than the bottom, provided a more consistent flat surface when baking at 65 $^{o}$C.
- It is very important that the soft PDMS substrates remain as clean as possible after creation; keeping them stored in a parafilm-closed petri dish and only opening the dish in a TC hood are useful for preventing dust buildup.

PVA film

- When making the PVA film, the PVA powder will only form a solution when heated. We preferred to heat the mixture in a 110 $^{o}$C oven for 30 minutes rather than using a microwave, as other literature suggests, in order to prevent boiling and loss of liquid. Filtering the liquid PVA was also useful for ensuring a homogeneous and flat film.
- Plasma-treating the petri dish before pouring in the liquid PVA ensured that the film remained attached to the petri dish and did not lift off prematurely.

Stamping

- When coating the PDMS stamp with fluorescent BSA, special care should be taken when rinsing the stamp with PBS. Remove the stamp from the PBS slowly and at an angle to ensure that as much liquid as possible is removed from the stamp. When PBS dries, it tends to leave behind salt crystals which can disrupt the pattern if it is allowed to dry on top of the stamp.
- If the concentration of BSA is too high, the pattern features may get filled in with fluorescence and result in a poor-looking pattern when transferred. Avoid too small features or too high concentration of BSA.
- When drying the stamp with Nitrogen gas after the final PBS rinse, use a low velocity stream as the fluorescent BSA layer can be fragile.
- When placing the PVA film onto the stamp, use a pair of tweezers to gently push on the film from the center outwards to eliminate bubbles between the two surfaces.

### 2. Methods

*PDMS stiffness measurements*

Dog bone-shaped samples of mixtures of PDMS (Sylgard 527 and 184) were created to measure their stiffness. Mixtures of 2.5%, 5%, and 10% ratios of Sylgard 184 to Sylgard 527 were poured into molds and cured at 65 $^{o}$C overnight. After curing, two spots used for measuring strain were drawn in the neck region of the sample. Several loads were hanged from a sample and imaged with a typical phone camera to evaluate the resulting engineering stress and strain for each load (Supplementary Figure 1A). Stress was calculated by dividing the applied weight by the minimum cross-sectional area of the sample (measured by analog calipers), and strain was calculated by measuring the distance between the two spots drawn previously. When plotted, the data points form a straight line where the slope of the line is the stiffness or Young's modulus. We measured stiffness on 3 separate samples for each PDMS mixture.

Our measurements found that mixtures containing 2.5%, 5%, and 10% Sylgard 184 results in stiffnesses or Young's modulus of 7.7, 13.5, and 46.7 kPa (Supplementary Figure 2B). These measurements agree closely with prior literature[29].

*Fibrinogen treatment of black dots*

Fibrinogen from human plasma that is conjugated to Alexa Fluor-488 (Life Technologies) was diluted to 0.15 mg/mL and incubated on the black dots for 1 hour. Substrates were mounted onto glass coverslips using Fluoromount-G mounting medium (Life Technologies) and imaged with confocal microscopy (Supplementary Figure 1H-J).

*Laminin treatment of black dots*

Mouse laminin (Life Technologies, 23017015) was diluted to 10 µg/mL and incubated directly onto black dots for 1 hour. Laminin was then labeled with a rabbit anti-laminin antibody (Millipore Sigma, L9393) and a goat anti-mouse IgG secondary antibody conjugated to Alexa Fluor 488 (Life Technologies, A-11008). The labeled substrate was then imaged by widefield microscopy (Supplementary Figure 1K-M).

### 3. Donor Specific Results

*Force-Area, Force-Circularity, and Force-F-actin Dispersion Relationships*

Supplementary Figures 3-5 show the Force-Area, Force-Circularity, and Force-F-actin dispersion results for each donor. Fit lines through the data were constrained to pass through the origin for Force-Area, but not for Force-Circularity or Force-F-actin dispersion. The slopes of these lines indicates the strength of the relationships, which is slightly different for each donor as shown in panel C of Supplementary Figures 3-5. An ANOCOVA with Tukey's post hoc test was used find significance between slopes.

*K-means Clustering*

Supplementary Figure 8 shows the K-means clustering performed for each donor separately. All 6 donors exhibited similar patterns when separated into two clusters, although donor 4 tends to cluster more strongly by area than other donors.

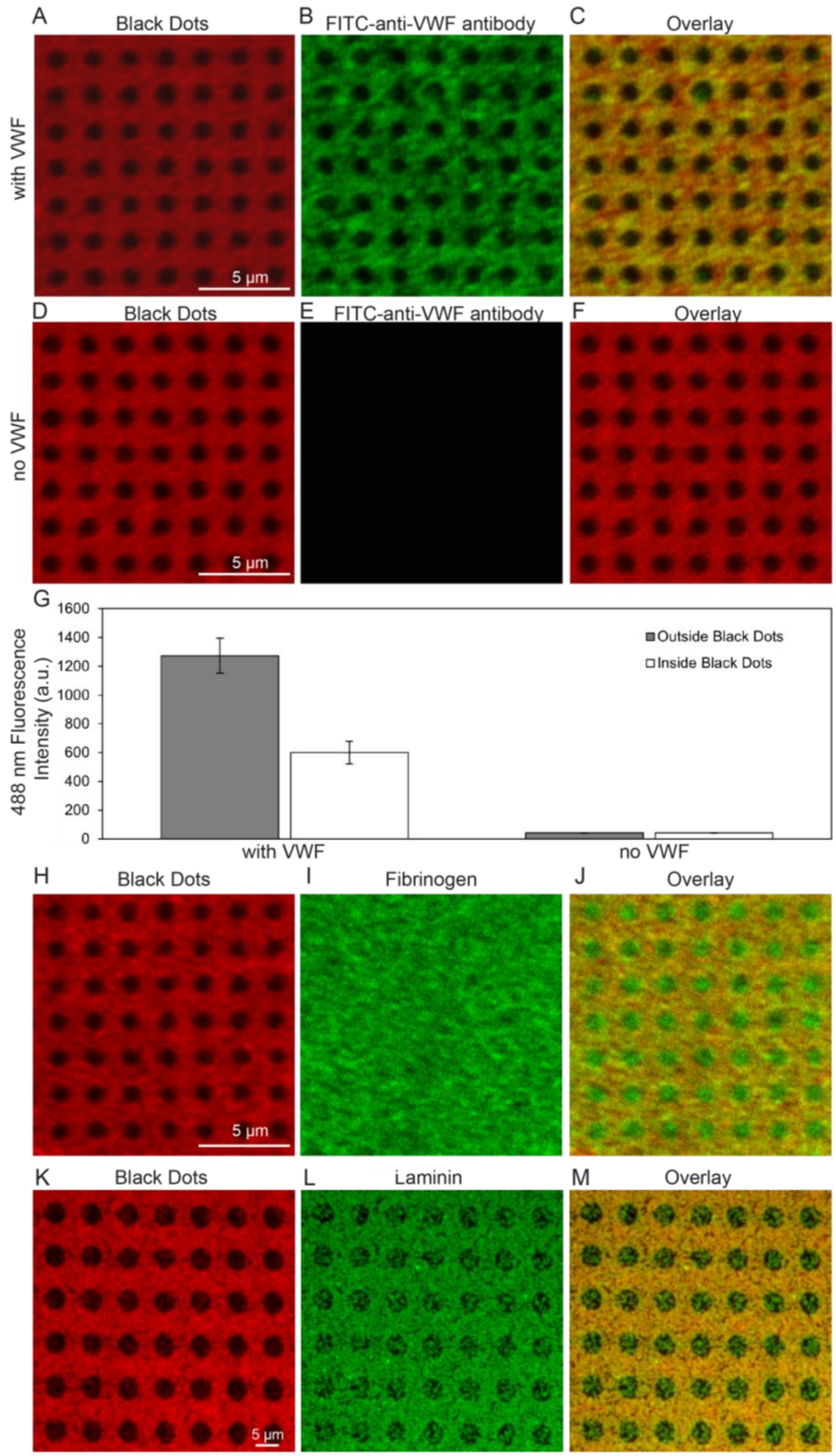


**Supplemental Figure 1 – ECM treatment of black dots.** (A-C) VWF-treated black dots with fluorescent BSA shown in red (A), VWF labeled with a FITC-anti-VWF antibody shown in green (B), and an overlay (C). Qualitatively, VWF is bound to both the fluorescent-BSA (orange in C) as well as the non-fluorescent black dots (green in C). (D-F) Black dots without VWF treatment with fluorescent BSA shown in red (D), VWF labeled with a FITC-anti-VWF antibody shown in green (E), and an overlay (F). As expected, the FITC-anti-VWF antibody does not bind to the substrate without VWF treatment (E). (G) Quantitatively, substrates with VWF have VWF bound both outside the black dots (gray bar) and inside the black dots (white bar), with higher fluorescent intensity outside the black dots. As expected, substrates without VWF have negligible 488 fluorescence both inside and outside the black dots. (H-I) Black dots can also be treated with fluorescent-fibrinogen, which binds uniformly across the surface, or (K-M) laminin, which has some preference for binding outside the black dots. Note that because we typically use laminin with larger cells (not platelets), its binding was characterized on black dots with a larger diameter. In this figure, the black dots are shown in red instead of orange because the red/green overlay is easier to interpret than the orange/green overlay.

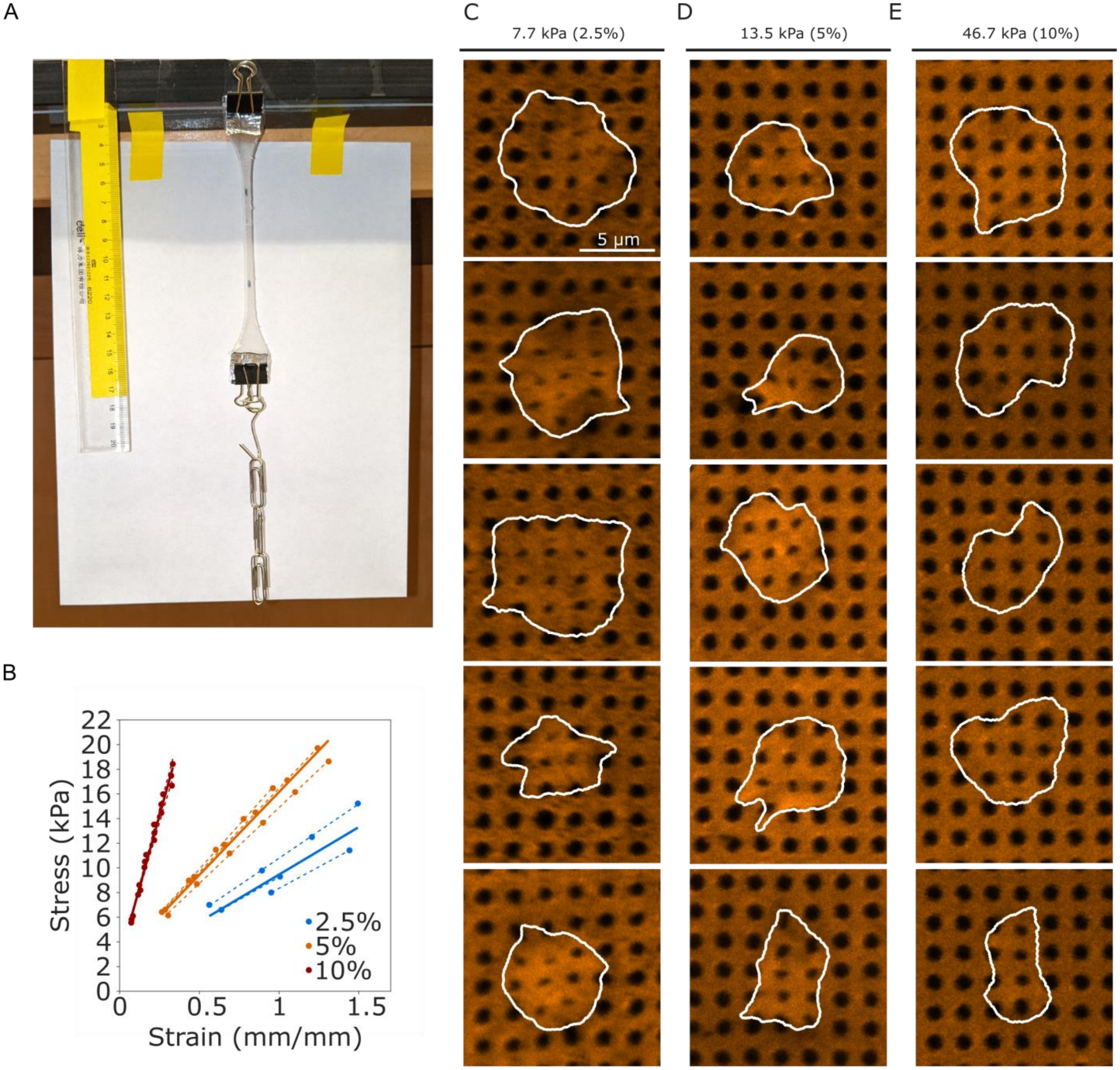


**Supplemental Figure 2 – Selection of substrate stiffness.** (A) PDMS tensile testing apparatus to measure the Young's modulus of each mixture of PDMS. (B) Measurements of stress and strain from the tensile test. Dotted lines indicate lines of best fit for each experiment. Solid lines are the average slope and intercept for the dashed lines for each given PDMS mixture. (C-E) Examples of platelets attached to black dots for the three different PDMS mixtures. Platelets on low stiffness (C) can distort the dots too much for image detection, while platelets on high stiffness (E) sometimes don't distort the dots beyond noise levels. The stiffness of 13.5 kPa in (D) was chosen for the studies presented in this paper.

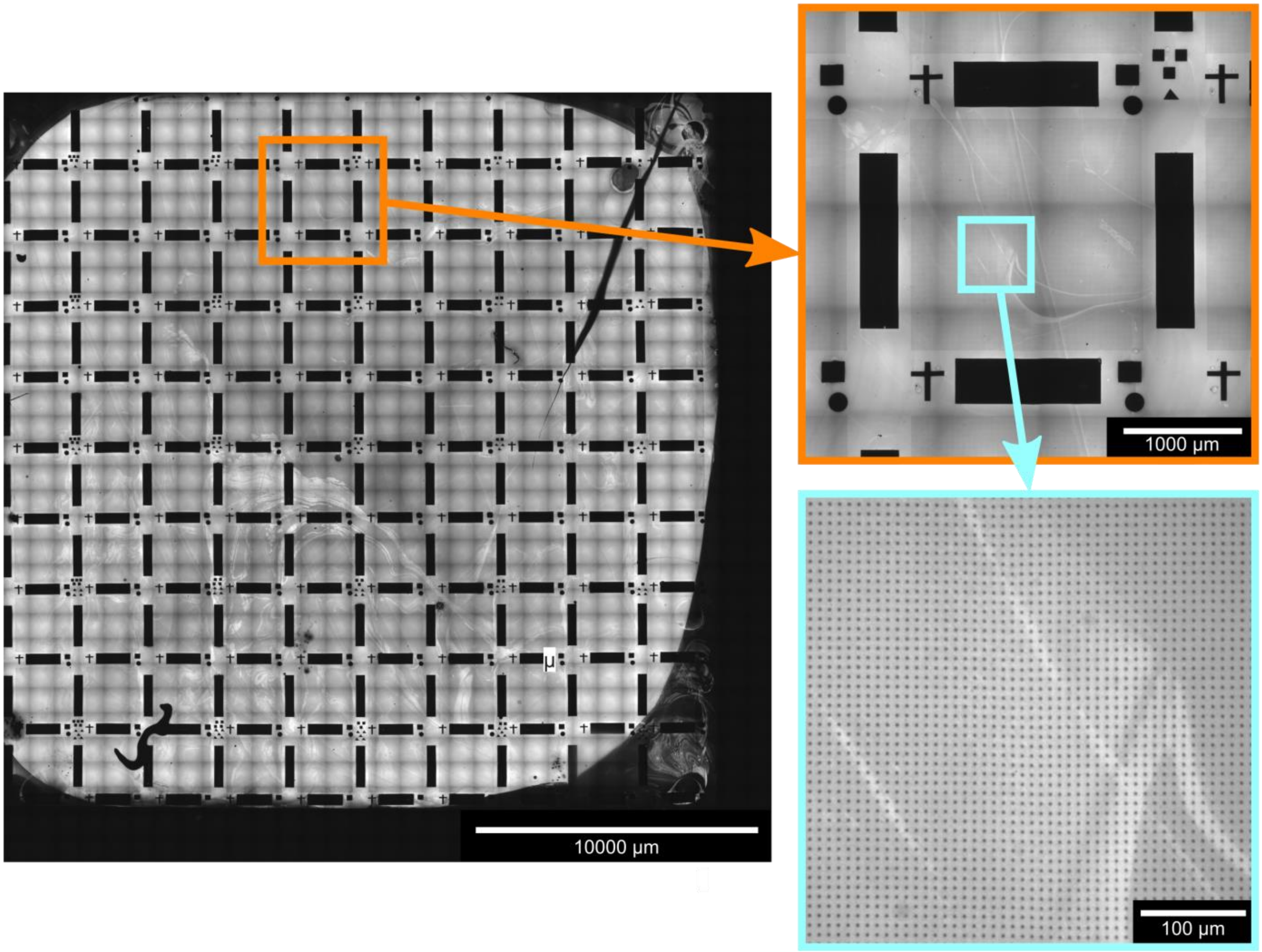


**Supplemental Figure 3 – Example of black dots printed over larger areas.** In this example, black dots have 9 um center-to-center spacing and cover a total physical area of nearly 4 $cm^2$.

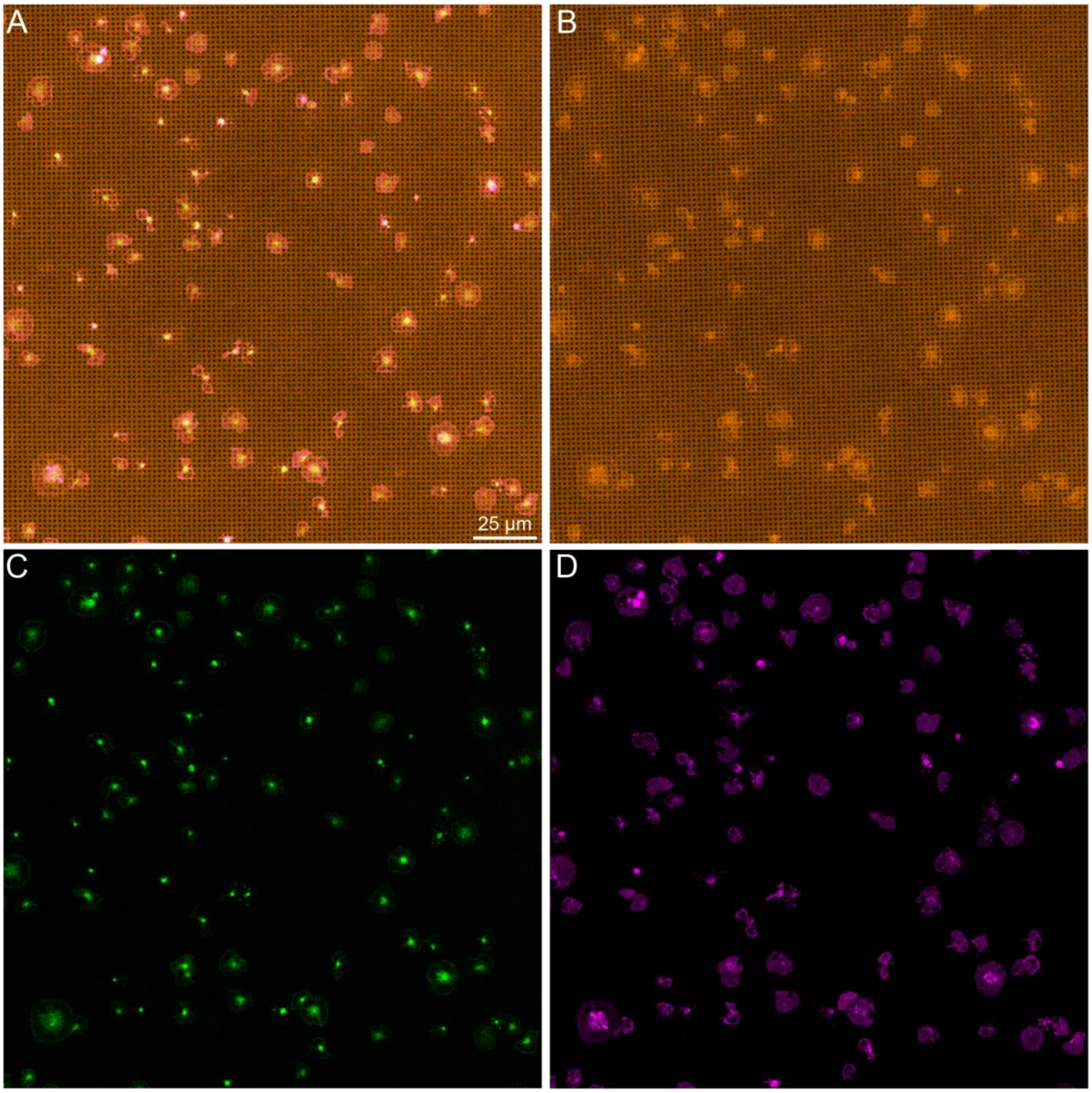


**Supplemental Figure 4 – Example of a field of view containing many platelets adhered to and contracting on the black dots.** This figure shows the same image as Figure 2A, but with each channel shown separately. Here, (A) is an overlay of all channels, (B) is the black dots in orange, (C) is F-actin in green, and (D) is GPIb in magenta.

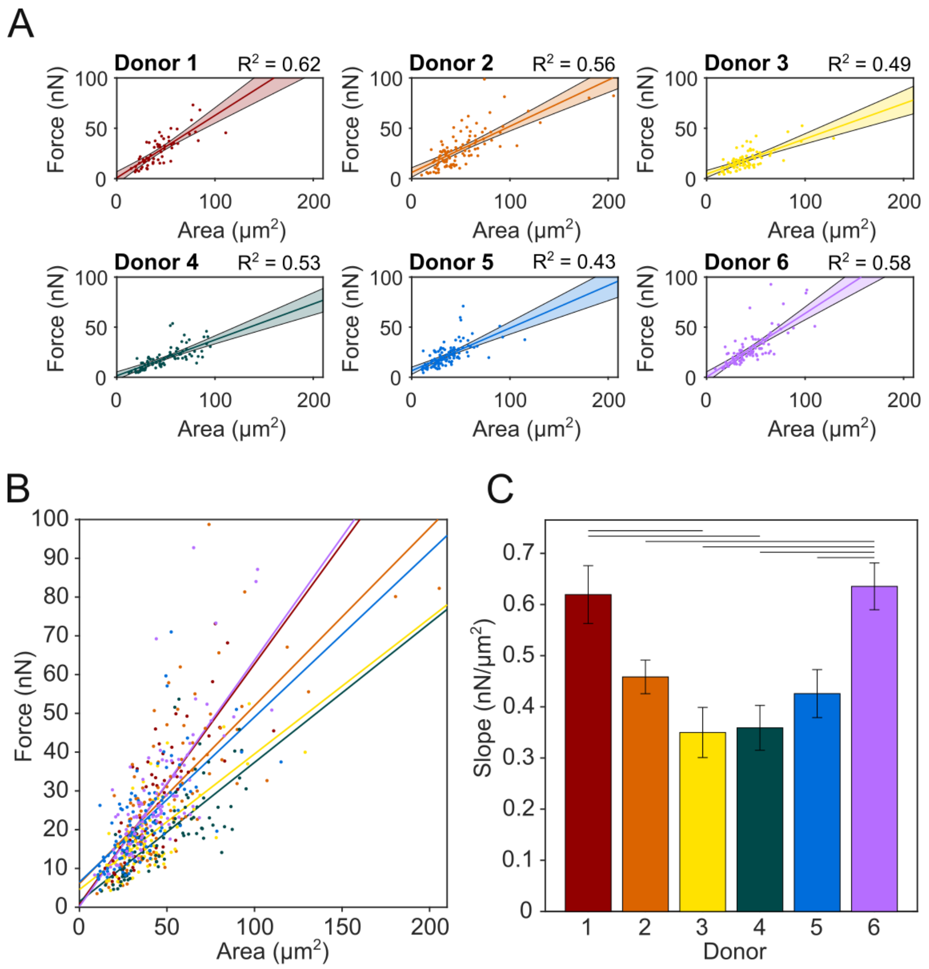


**Supplemental Figure 5 – Force-area relationship by donor.** (A) Force-area data plotted separately for each donor. Solid fit lines are displayed with shaded area that indicates 95% prediction interval for the fits. (B) Overlay of force-area data from all 6 donors with linear fit lines for each donor. (C) Slopes for the fit lines for each donor. Horizontal bars at the top indicate significance ($p < 0.05$ when tested with a one-way ANOCOVA and Tukey's post hoc test).

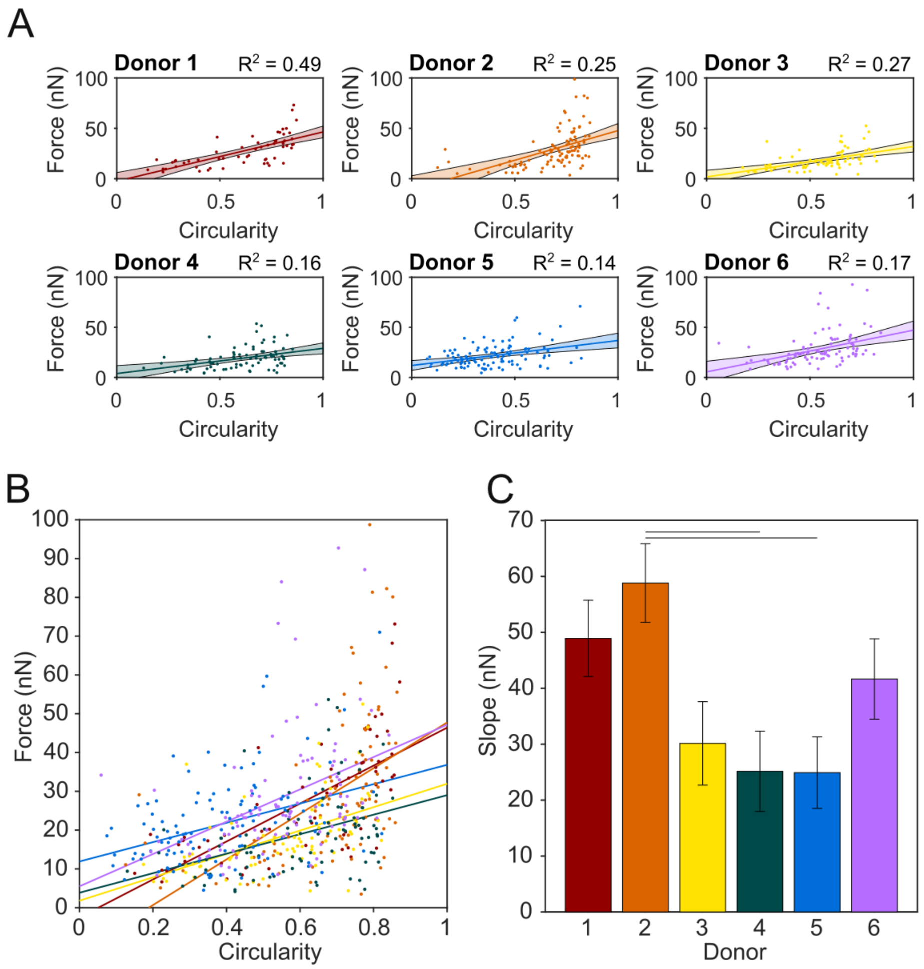


**Supplemental Figure 6 – Force-circularity relationship by donor.** (A) Force-circularity data plotted separately for each donor. Solid fit lines are displayed with shaded area that indicates 95% prediction interval for the fits. (B) Overlay of force-circularity data from all 6 donors with linear fit lines for each donor. (C) Slopes for the fit lines for each donor. Horizontal bars at the top indicate significance ($p < 0.05$ when tested with a one-way ANOCOVA and Tukey's post hoc test).

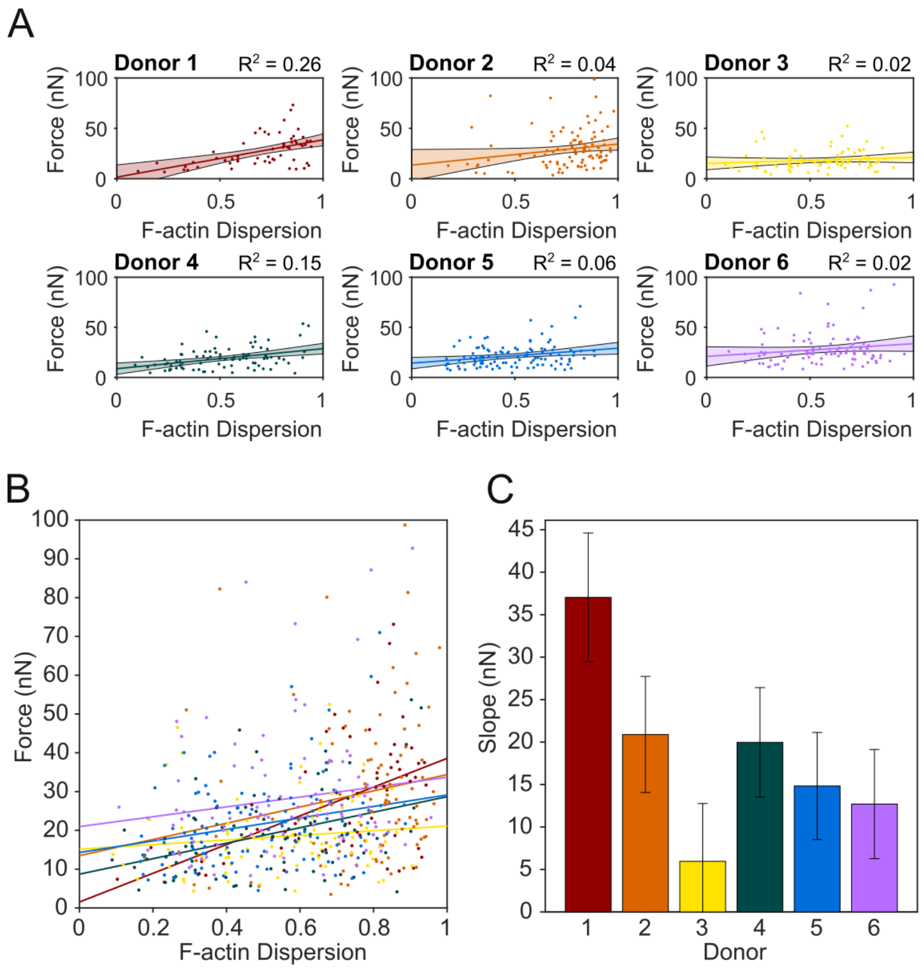


**Supplemental Figure 7 – Force-F-actin dispersion relationship by donor.** (A) Force-area data plotted separately for each donor. Solid fit lines are displayed with shaded area that indicates 95% prediction interval for the fits. (B) Overlay of force-area data from all 6 donors with linear fit lines for each donor. (C) Slopes for the fit lines for each donor. No significance was detected for the slopes between donors when tested with a one-way ANOCOVA and Tukey's post hoc test.

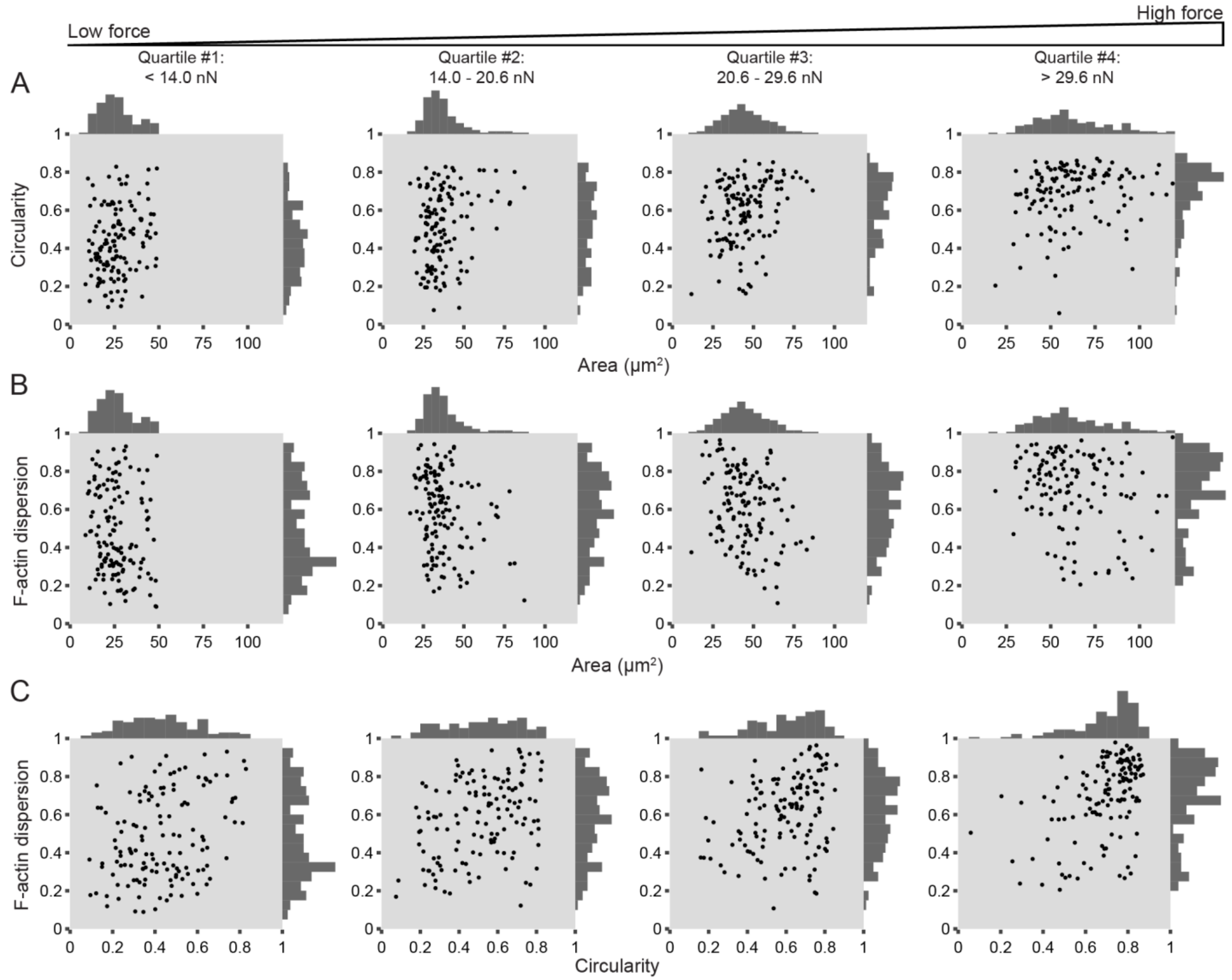


**Supplemental Figure 8 – Platelet size, shape, and structure do not strongly correlate with each other, but increase together with force.** This figure shows the same data as main text Figure 4B, E, and I, but with a scatter plot instead of a contour plot to display all points of data.

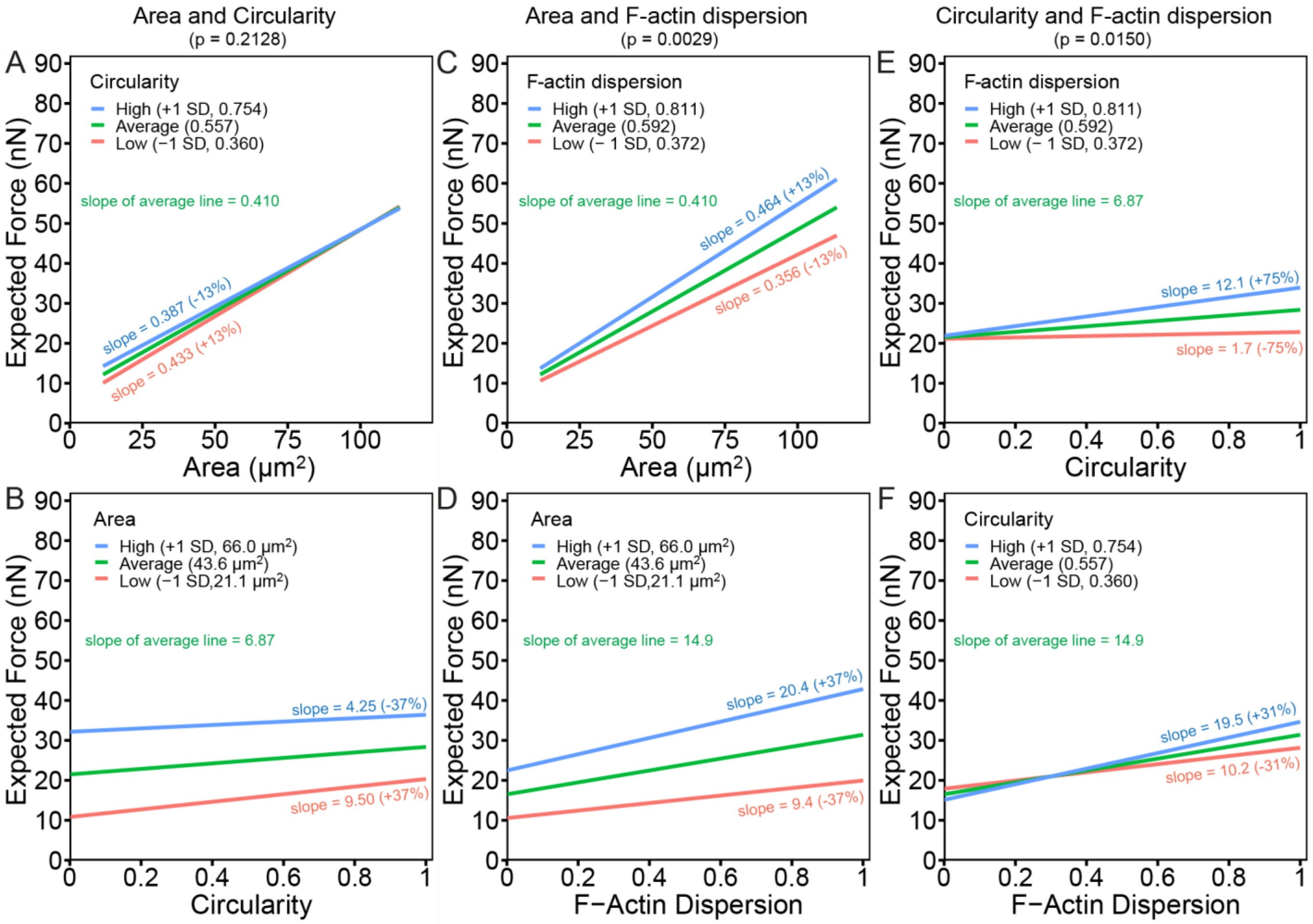


**Supplemental Figure 9 – Interaction plots demonstrate cooperative interactions between F-actin dispersion and circularity as well as F-actin dispersion and area.** These plots are output from the multivariate mixed effects model. They demonstrate the relationship between expected force (y-axis) and the x-axis variable change when another variable is average (green line), high (blue line), or low (red line). Slope differences in these green, blue, and red lines indicate how one variable changing alters the relationship between force and the x-axis variable. (A) For average circularity (circularity = 0.557, green line), the relationship between expected force and area is positive with a slope of 0.410 nN/μm$^2$. If circularity is one standard deviation above average (circularity = 0.754, blue line), the force-area relationship has a slope of 0.387 nN/μm$^2$. If circularity is one standard deviation below average (circularity = 0.360, red line), the force-area relationship has a slope of 0.433 nN/μm$^2$. The interaction between area and circularity is not significant (p = 0.2128) (B) For average area (area = 43.6 μm$^2$, green line), the relationship between expected force and circularity is positive with a slope of 6.87 nN/1 circularity unit. If area is one standard deviation above average (area = 66.0 μm$^2$, blue line), the force-circularity relationship has a slope of 4.25 nN/1 circularity unit. If area is one standard deviation below average (area = 21.1 μm$^2$, red line), the force-circularity relationship has a slope of 9.50 nN/1 circularity unit. The interaction between area and circularity is not significant (p = 0.2128) (C) For average F-actin dispersion (F-actin dispersion = 0.592, blue line), the relationship between expected force and area is positive with a slope of 0.410 nN/μm$^2$. If F-actin dispersion is one standard deviation above average (F-actin dispersion = 0.811, blue line), the force-area relationship has a slope of 0.464 nN/μm$^2$. If F-actin dispersion is one standard deviation below average (circularity = 0.372, red line), the force-area relationship has a slope of 0.356 nN/μm$^2$. The interaction between area and F-actin dispersion is significant (p = 0.0029). (D) For average area (area = 43.6 μm$^2$, green line), the relationship between expected force and F-actin dispersion is positive with a slope of 14.9 nN/1 F-actin dispersion unit. If area is one standard deviation above average (area = 66.0 μm$^2$, blue line), the force-F-actin dispersion relationship has a slope of 20.4 nN/1 circularity unit. If area is one standard deviation below average (area = 21.1 μm$^2$, red line), the force-F-actin dispersion relationship has a slope of 9.4 nN/1 F-actin dispersion unit. This interaction between area and F-actin dispersion is significant (p = 0.0029). (E) For average F-actin dispersion (F-actin dispersion = 0.592, blue line), the relationship between expected force and circularity is positive with a slope of 6.87 nN/1 circularity unit. If F-actin dispersion is one standard deviation above average (F-actin dispersion = 0.811, blue line), the force-circularity relationship has a slope of 12.1 nN/1 circularity unit. If F-actin dispersion is one standard deviation below average (circularity = 0.372, red line), the force-circularity relationship has a slope of 1.7 nN/1 circularity unit The interaction between circularity and F-actin dispersion is significant (p = 0.0150). (F) For average circularity (circularity = 0.557, green line), the relationship between expected force and F-actin dispersion is positive with a slope of 14.9 nN/1 F-actin dispersion unit. If circularity is one standard deviation above average (circularity = 0.754, blue line), the force-F-actin dispersion relationship has a slope of 19.5 nN/1 unity of F-actin dispersion. If circularity is one standard deviation below average (circularity = 0.360, red line), the force-F-actin dispersion relationship has a slope of 10.2 nN/1 unit of F-actin dispersion. The interaction between circularity and F-actin dispersion is significant (p = 0.0150).

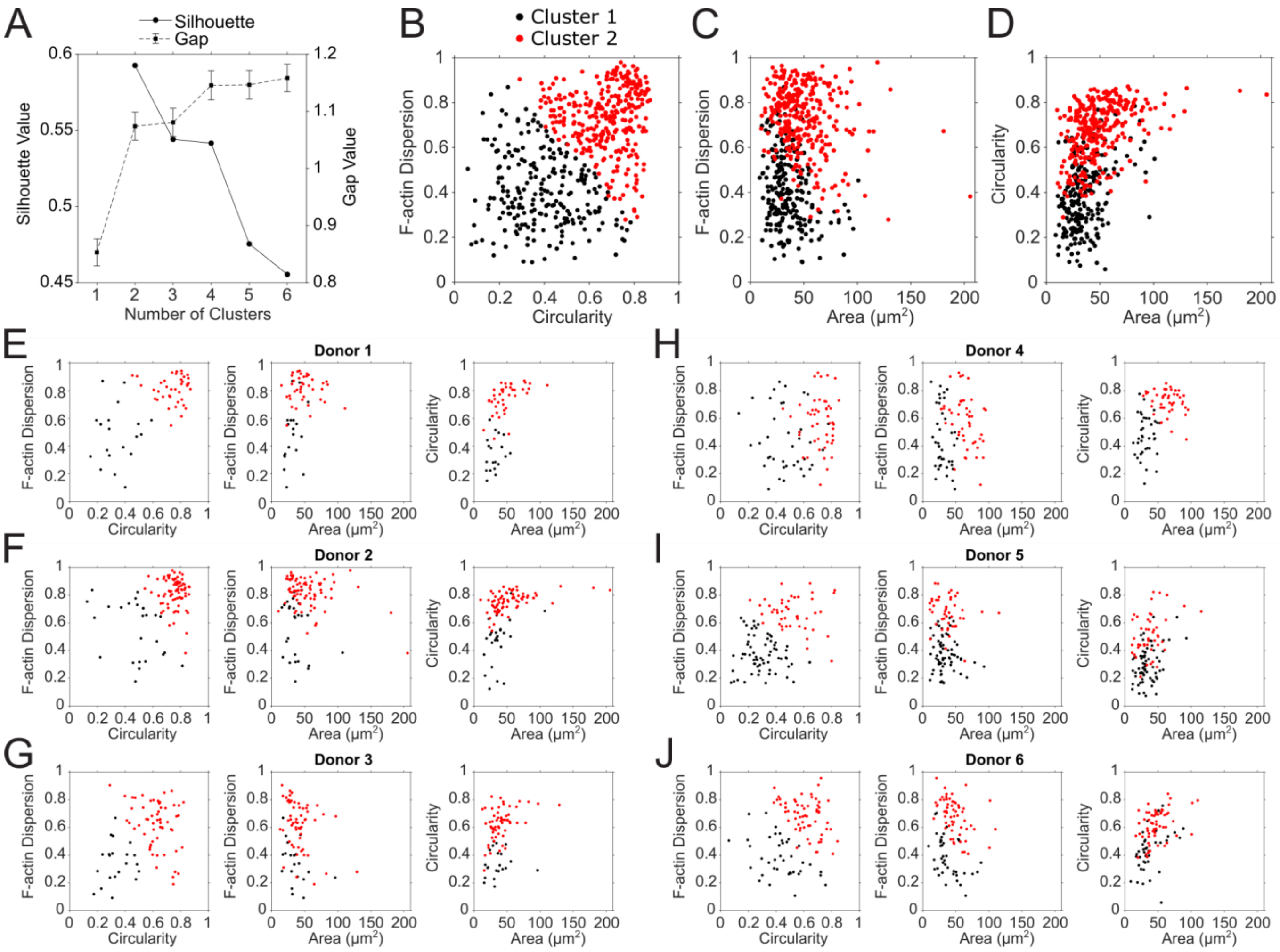


**Supplemental Figure 10 – K-means clustering and for each donor.** (A) Silhouette and gap values used to determine the optimal number of clusters. For silhouette, the highest value determines the optimal number of clusters. For gap, the optimal number of clusters is the lowest number where the gap value of the next highest number is within the error of the previous one. In this case, both methods indicate that 2 clusters is optimal. (B, C, D) Same data as shown in the main text Figure 5A, B, and C, except the area axis shows all data points. (E-J) Relationships between area, circularity, and F-actin dispersion for all 6 donors. Here, K-means clustering was performed on each donor separately and generally shows that the platelets separate into two similar clusters for each donor.
